# Structural Basis for an Unprecedented Enzymatic Alkylation in Cylindrocyclophane Biosynthesis

**DOI:** 10.1101/2021.12.02.470901

**Authors:** Nathaniel R. Braffman, Terry B. Ruskoski, Katherine M. Davis, Nate Glasser, Cassidy Johnson, C. Denise Okafor, Amie K. Boal, Emily P. Balskus

## Abstract

The cyanobacterial enzyme CylK assembles the cylindrocyclophane natural products by performing two unusual alkylation reactions, forming new carbon-carbon bonds between aromatic rings and secondary alkyl halide substrates. This transformation is unprecedented in biology and the structure and mechanism of CylK are unknown. Here, we report x-ray crystal structures of CylK, revealing a distinctive fusion of a Ca^2+^ binding domain and a β-propeller fold. We use a mutagenic screening approach to locate CylK’s active site at its domain interface, identifying two residues, Arg105 and Tyr473, that are required for catalysis. Anomalous diffraction datasets collected with bound bromide ions, a product analog, suggest these residues interact with the alkyl halide electrophile. Additional mutagenesis and molecular dynamics simulations implicates Asp440 and Glu374 in activating the nucleophilic aromatic ring. Bioinformatic analysis of CylK homologs from other cyanobacteria establishes that they conserve these key catalytic amino acids but they are likely associated with divergent reactivity and altered secondary metabolism. By gaining a molecular understanding of this unusual biosynthetic transformation, this work fills a gap in our understanding of how alkyl halides are activated and used by enzymes as biosynthetic intermediates, informing enzyme engineering, catalyst design, and natural product discovery.

## Introduction

Chemists utilize alkyl halides (molecules with sp^3^ C–X bonds, X = F, Cl, Br, or I) as key synthetic reagents because of their accessibility and favorable reactivity (1); however, examples of their use as intermediates in biological systems are rare (2). A few characterized natural product biosynthetic pathways generate transiently halogenated intermediates that facilitate downstream chemical reactions in a strategy known as ‘cryptic halogenation’ (3). This biosynthetic logic requires two partner enzymes: a halogenase, which introduces the alkyl halide substituent, and a second enzyme that utilizes the halogenated intermediate as a substrate, leveraging its increased reactivity and releasing the corresponding halide anion as a side product. Multiple classes of halogenases have been structurally and mechanistically characterized (4), but the cognate halide-utilizing enzymes in pathways that employ cryptic halogenation remain underexplored. In particular, the structures and biochemical mechanisms for enzymatically engaging and activating alkyl halide substrates as intermediates in natural product biosynthesis are unknown.

Interrogating cylindrocyclophane biosynthesis by the cyanobacterium *Cylindrospermum licheniforme* ATCC 29412 presents an exciting opportunity to study alkyl halide utilization in biological systems. In this organism, the putative diiron carboxylate halogenase CylC generates an alkyl chloride intermediate which is further elaborated to produce resorcinol-containing alkyl chloride **1**. Two molecules of **1** are then dimerized by an alkyl chloride-utilizing enzyme, CylK, via the formation of two new carbon-carbon (C–C) bonds to construct a paracyclophane ring system (Fig. 1A) (5). CylK is annotated as a fusion of a Ca^2+^ binding domain and a β-propeller fold, but the roles of these predicted protein domains in catalysis are unknown. Initial biochemical studies revealed that this transformation involves two stereoselective alkylation events that occur in a stepwise fashion, with inversion of configuration at the alkyl chloride stereocenter. CylK is the only enzyme known to catalyze aromatic ring alkylation with an alkyl halide electrophile, a reaction that mirrors a classical non-enzymatic reaction known as the Friedel–Crafts alkylation, which is an important transformation in organic synthesis. The traditional non-enzymatic Friedel– Crafts reaction suffers from a lack of stereo- and regiocontrol, unwanted overalkylation events, and the generation of carbocationic intermediates that can undergo unproductive rearrangements (6, 7).

**Figure 1.**
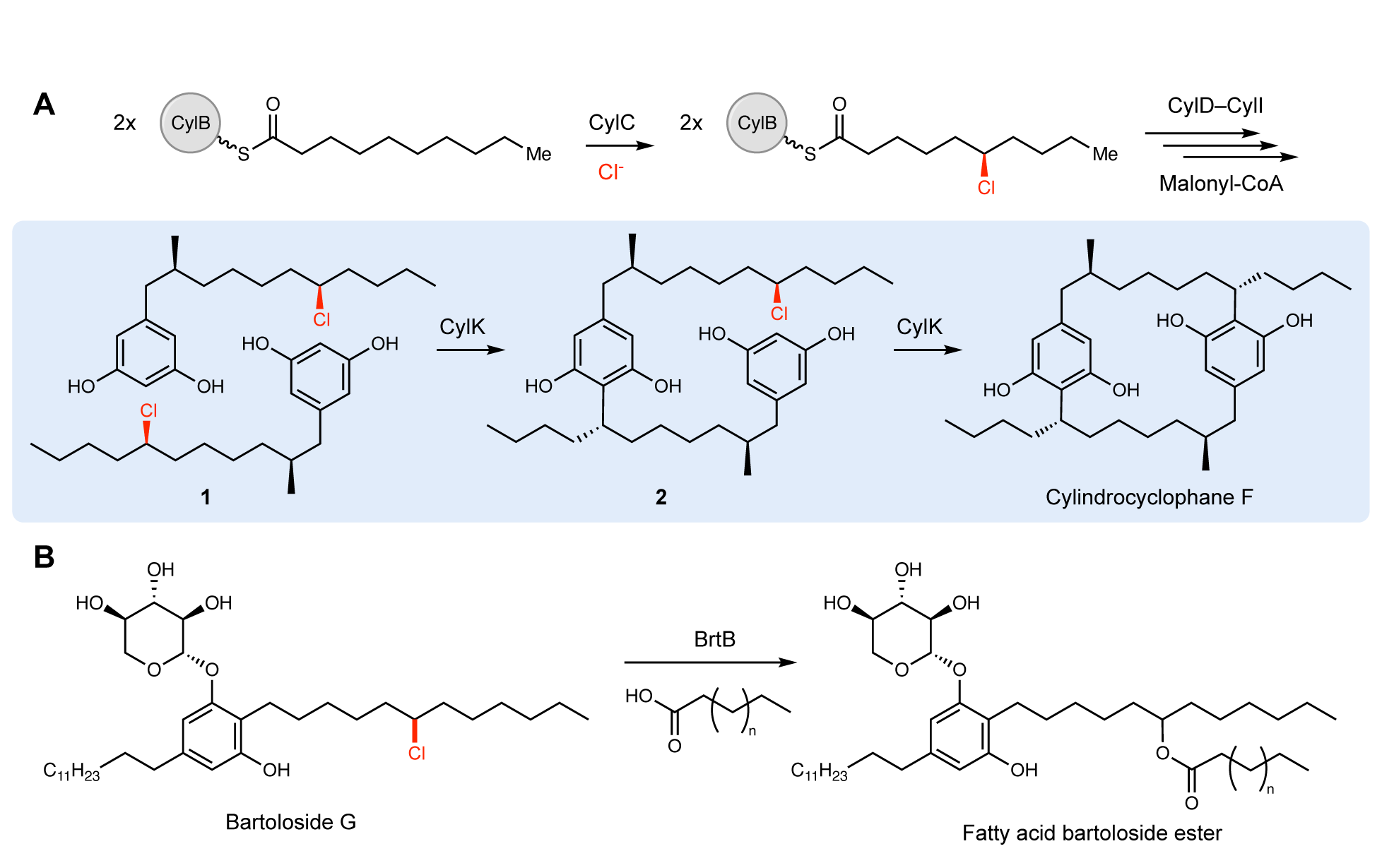
CylK and related enzymes use alkyl chloride substrates as biosynthetic intermediates. (*A*) The halogenase CylC generates an alkyl chloride substrate for CylK, which catalyzes two stepwise Friedel– Crafts alkylations to construct a paracyclophane macrocycle in cylindrocyclophane biosynthesis. This involves an intermolecular reaction between two resorcinol-containing alkyl chloride substrates (**1**) to generate intermediate **2**, followed by an intramolecular alkylation to afford cylindrocyclophane F, shaded in blue. (*B*) The related enzyme BrtB (30% amino acid identity, 46% similarity) catalyzes an analogous but chemically distinct C–O bond-forming event between chlorinated bartolosides and fatty acid nucleophiles.

In contrast, the CylK-catalyzed Friedel–Crafts alkylation overcomes these limitations, likely by tightly controlling reactivity within the enzyme active site. This enzymatic transformation holds promise for biocatalytic applications, and we have recently demonstrated CylK’s ability to accept multiple discrete resorcinol nucleophiles and alkyl halide electrophiles with varying substitution and electronic character (8). Moreover, uncharacterized enzymes with homology to CylK are found in numerous cyanobacteria, one of which (BrtB) was recently revealed to catalyze carbon-oxygen (C–O) bond formation between the carboxylate groups of fatty acids and bartoloside A, an alkyl chloride-containing natural product (Fig. 1B) (9). This suggests that the mechanism by which CylK binds and activates alkyl chlorides might be shared with this and other homologs that use diverse nucleophilic substrates.

Despite the intriguing reactivity and potential applications of CylK and related alkyl halide activating enzymes, our knowledge of its structure and mechanism is limited. In this work, we present a crystal structure of CylK, identify critical active site residues, provide experimental and bioinformatic support for their roles in catalysis, and propose a mechanism by which alkyl chloride substrates are activated for stereospecific nucleophilic substitution. In determining how CylK performs this Friedel–Crafts alkylation, we have enhanced our fundamental understanding of how enzymes engage alkyl halide substrates. Our structural information and mechanistic model will guide future enzyme engineering efforts, inform the design of non-enzymatic catalysts, and enable genome mining to uncover new natural products constructed by related biosynthetic strategies.

## Results

### CylK is a distinctive fusion of two protein domains

CylK crystallized in the *C*222_1_ space group with a single monomer in the asymmetric unit (Table S1). Suitable molecular replacement (MR) models for phasing could not be identified. Phase information was obtained by partial substitution of native Ca^2+^ binding sites in CylK with Tb^3+^, followed by collection of X-ray diffraction datasets at the Tb X-ray absorption peak energy. The structure of CylK was solved to 1.68 Å resolution via a combined MR and single-wavelength anomalous diffraction (SAD) approach, with the MR search model generated from a poly-alanine 7-bladed β-propeller. The structure reveals an unprecedented fusion of two protein folds, an N-terminal Ca^2+^ binding domain and a C-terminal β-propeller domain (Fig. 2A).

**Figure 2.**
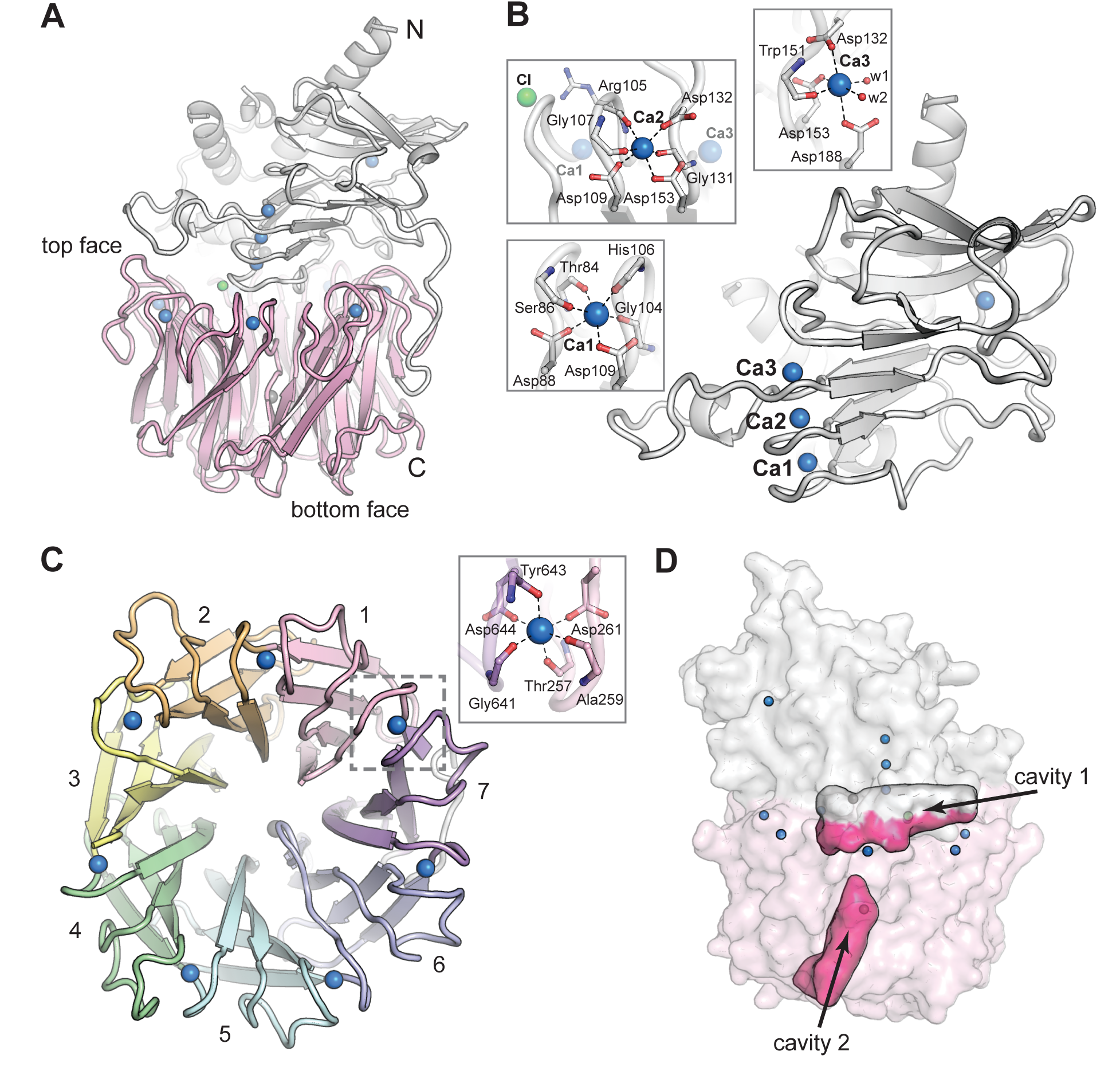
The CylK X-ray crystal structure reveals a distinct arrangement of two protein domains. (*A*) An overall view of the structure of CylK. In this image and throughout, calcium ions are shown as blue spheres, magnesium ions as dark grey spheres, and chloride ions as green spheres. The N-terminal domain is depicted as a light grey ribbon diagram and the C-terminal domain is shown as a pink ribbon diagram. (*B*) The N-terminal domain contains a right-handed parallel β-roll stabilized by three Ca^2+^ ions. The structure is capped by a three-strand antiparallel β-sheet and buttressed by additional helical secondary structures. Insets show the Ca^2+^ ion coordination environment within the β-roll. (*C*) Cavity mapping analysis (56) of CylK reveals two cavities large enough to accommodate the alkyl resorcinol substrates. (*D*) A top-down view of the 7-bladed β-propeller C-terminal domain. The propeller blades are numbered from the N-terminal end of the domain and colored by blade. The inset shows a representative view of the Ca^2+^ coordination environment. A similar binding site exists at each blade junction.

The N-terminal domain of CylK contains a short helical component that packs against a β-roll core (Fig. 2B & S1). The core fold is structurally similar to repeat-in-toxin (RTX) motifs, found in diverse bacterial extracellular proteins (10). A search of the PDB for structural relatives of the CylK N-terminal domain reveals similarity to the C-terminal RTX domains of secreted toxins (11), such as the adenylate cyclase toxin of *Bordatella pertussis*, and bacterial surface-layer proteins (12), such as the RsaA protein of *Caulobacter crescentus* (Table S2). In these systems, the RTX domain facilitates Ca^2+^-dependent folding or assembly in the calcium-rich environment outside the cell but not in the calcium-depleted cytosol. The CylK N-terminal domain also resembles RTX domains fused to catalytic domains of extracellular hydrolases and epimerases from gram-negative bacteria (Fig. S2) (13–15). In these systems, the RTX unit is attached to the C-terminus of the catalytic domain to facilitate extracellular secretion and Ca^2+^-dependent folding. The RTX component of these characterized proteins is functionally modular, with no obvious direct connection to the active site, although in some systems that target large biopolymer substrates, the RTX domain may help anchor the substrate via electrostatic interactions (14).

The N-terminal domain of CylK differs from typical RTX motifs in several important ways. Canonical RTX motifs have an extended compact oval structure shaped by tight turns between subsequent β-strands that are stabilized by stacked metal-binding motifs on both sides of the core fold (10). The metal binding sites are formed by a repeating GXXGXD motif in the tight turn. Vertically stacked pairs of these repeats yield hexacoordinate Ca^2+^ binding sites with four carbonyl ligands and two carboxylates provided by the conserved Asp side chains. CylK adopts a more asymmetric version of this fold and contains just a single copy of the consensus Ca^2+^ binding motif. The modified β-roll structure in CylK contains three vertically aligned Ca^2+^ ions on one side of the core fold. The other side lacks the metal binding sites found in other RTX proteins. Additionally, while the first two turns of the β-roll in CylK form tight junctions near the Ca^2+^ ions, all of the subsequent turns contain long extensions (Fig. S1), some of which facilitate integration with the C-terminal domain. Interestingly, the section of the β-roll motif that conserves the Ca^2+^ sites packs closely into the center of the C-terminal domain. Although the CylK N-terminal domain is topologically distinct from other β-roll motifs, CylK shares with these systems a functional requirement for Ca^2+^ (5).

The C-terminal domain of CylK adopts a 7-bladed β-propeller fold (Fig. 2C & S3), a widespread structural motif found in both eukaryotic and prokaryotic protein structures (16). In eukaryotes, this flat, cylindrically shaped fold facilitates protein-protein interactions (17) via the same interface (top face) that interacts with the N-terminal domain in CylK. In prokaryotes and plants, β-propeller folds can be used in catalysis (16). Comparison of the CylK β-propeller domain to other examples of this fold in bacterial enzymes reveals several structural differences unique to CylK (Fig. S4). In all β-propellers, adjacent four-stranded “blades” are connected back-to-front by loops on the top face of the domain. The top face of the fold also consists of loops that link the internal β-strands of each propeller blade. In CylK, these internal loops are unusually long and folded back over the outside of the central propeller fold. Also, the loops that connect subsequent blades contain unique blade-bridging Ca^2+^ binding sites, suggesting a shared role for this divalent metal in structural stabilization of both domains of CylK.

CylK is structurally similar to a class of single domain bacterial β-propeller enzymes implicated in streptogramin antibiotic resistance that includes virginiamycin B lyase (Vgb) (Table S3) (18, 19). These enzymes share with CylK an ability to interface with macrocyclic substrates or products, but they perform a distinct chemical transformation. While CylK forms two C–C bonds to generate a cyclic product, the streptogramin resistance proteins linearize cyclic peptide substrates by cleaving a C–O bond. X-ray crystal structures of an inactive variant of Vgb bound to a substrate analog provide insight into the location of the active site and mechanism of macrocycle opening (18). In Vgb, the substrate analog and a Mg^2+^ ion necessary for catalysis bind near conserved polar side chains in blades 6 and 7 of the propeller fold (Fig. S4B). In CylK, these sites are substituted for hydrophobic side chains, suggesting that, despite the similarities to streptogramin lyases, the substrates of CylK likely bind in a different location and are transformed by a distinct mechanism.

### The CylK active site is located at the domain interface

To identify CylK’s active site, we focused on two solvent accessible cavities (Fig. 2D). One cavity is lined by the top face of the β-propeller domain and by the N-terminal domain, forming a solvent accessible tunnel that is ∼18 Å deep and 15 Å wide at its largest point. The walls of this cavity contain both charged/polar residues and hydrophobic patches, ideal for interaction with the amphipathic resorcinol substrates (Fig. S5). We also interrogated a second central channel located exclusively within the C-terminal domain and opening to the opposite (bottom face) side of the propeller motif. To identify the active site, we individually mutagenized 17 polar residues spanning the two cavities (Fig. 3A) that are conserved in putative CylK homologs from cyanobacteria known to produce structurally related natural products (Fig. S6) (20–23). To rapidly test variant proteins for activity, we designed a plate-based lysate activity assay using non-native substrate pair **3** + **4** (Fig. 3B), which was previously demonstrated to be accepted by CylK (8). All variants associated with the central/bottom channel of the C-terminal β-propeller retained full alkylation activity (Fig. 3C). Strikingly, substitution of two residues located at the domain interface between the top face of the β-propeller and the N-terminal domain, Arg105 and Tyr473, completely abolished all activity on substrate pair **3** + **4** (Fig. 3C). Notably, although located in close proximity, Arg105 is supplied by the N-terminal domain, while Tyr473 is from the C-terminal β- propeller, suggesting that both domains play an essential role in catalysis. These data preliminarily supported the location of the active site as the cleft formed between the two domains of CylK.

**Figure 3.**
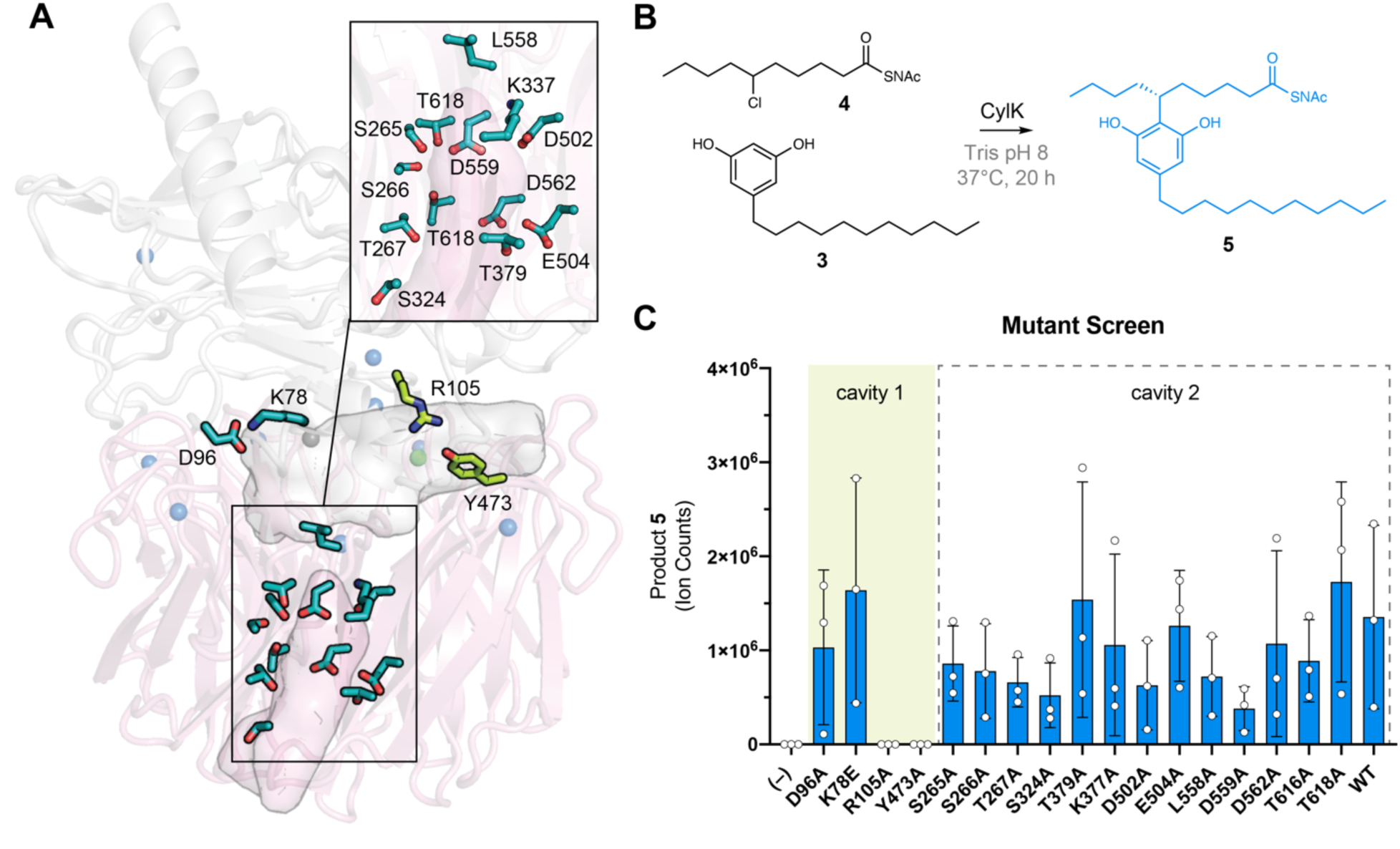
The active site of CylK is located at the domain interface. (*A*) Selected amino acids in CylK near cavities 1 and 2 subjected to mutant scanning are shown in stick format. Side chains shown in yellow near cavity 1 correspond to sites found to be essential for activity. (*B*) Non-native substrate pair used in CylK activity assays and product of the alkylation reaction (SNAc = *N*-Acetylcysteamine). (*C*) Screen of mutant activity to locate the CylK active site. Product formation was measured by LC-MS (Liquid Chromatography-Mass Spectrometry), error bars represent the standard deviation from the mean of three biological replicates.

In parallel, we hypothesized that the alkyl chloride moiety may be an important substrate-binding determinant based on our inability to obtain a full occupancy complex of resorcinol **3** in CylK crystals. Although **3** was required for CylK crystallization, we could not model the alkyl resorcinol substrate in the resulting structure. To visualize locations within CylK capable of binding an alkyl halide or free halide, we soaked CylK crystals with NaBr solutions. The Br^−^ ions are surrogates for the native Cl^−^ side product and are generated in reactions with alkyl bromide electrophiles, which CylK was previously shown to accept with similar efficiency to alkyl chlorides (8). Bromine X-ray absorption energies can be easily accessed at a conventional synchrotron X-ray source for anomalous diffraction experiments, allowing us to pinpoint the location of the halide in the structure. Initial refinements identified two strong positive peaks located ∼15 Å apart in *F*_o_−*F*_c_ electron density maps of the interdomain cavity. Anomalous diffraction datasets collected at the absorption edge of bromine confirmed assignment of these peaks as Br^−^ ions (Fig. 4A & Table S4). Anomalous peaks were not found in the central channel of the β-propeller domain.

**Figure 4.**
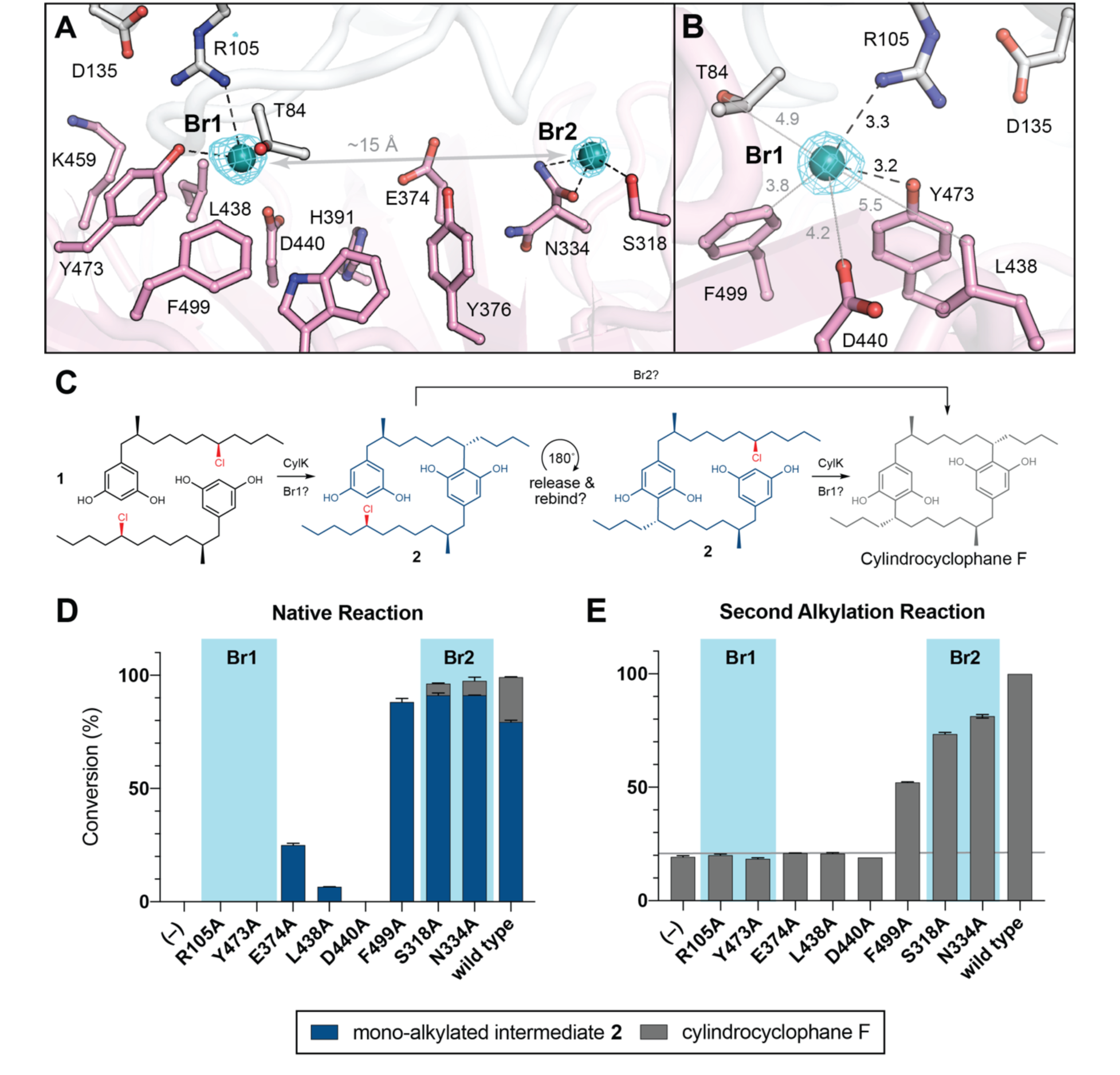
NaBr soak and mutagenesis of bromide-binding residues suggest a single site for catalysis. (*A*) A soak of NaBr into CylK crystals reveals two strong peaks in the anomalous difference electron density map (cyan mesh, contoured at 5.0 σ) within cavity 1. Bromide ion 1 (Br1) and 2 (Br2) are shown as teal spheres and selected amino acids in the vicinity of each site are shown in stick format. The C-terminal domain is colored white and the N-terminal domain is colored pink. (*B*) Alternate view of Br1 with anomalous difference electron density map, polar contacts shown as black dashed lines, other distances shown as grey lines. All distances measured in angstroms. (*C*) Potential roles for the two bromide binding sites in catalysis of paracyclophane formation. (*D*) End-point activity at 1 hour of select mutants performing the native reaction with substrate **1**, highlighting the residues associated with Br1 or Br2. Product formation was quantified by HPLC, error bars represent the standard deviation from the mean of two biological replicates. (*E*) End-point activity at 22 hours of select mutants performing the second alkylation reaction with intermediate **2**.

The bromide peak of highest intensity (∼33 σ) in the anomalous difference electron density map, Br1, is located just inside the opening to the interdomain cavity (Fig. 4B). Br1 is modeled at nearly full (85%) occupancy and appears to make hydrogen bonding contacts with the side chains of Arg105 from the N-terminal domain and Tyr473 from the C-terminal domain, both residues identified in our mutagenesis scan. Notably, the apo CylK model also contained a positive *F*_o_−*F*_c_ electron density peak at the Br1 site. Given the presence of chloride salts in the protein storage buffer, we modeled this site as a chloride ion in the apo model (Fig. 2B). Br1 resides within 8 Å of the calcium-binding sites in the N-terminal domain, and the backbone carbonyl of Arg105 coordinates the central Ca^2+^ ion (Ca2) in the β-roll motif of the N-terminal domain. This observation further supports an active role for both domains in the CylK catalyzed reaction. Br1 is located within 5-6 Å of other side chains that form a portion of the solvent accessible channel nearest to the protein surface (Fig. 4B), including a number of polar residues that could be implicated in activation of the resorcinol nucleophile and/or the alkyl chloride electrophile. Two residues, Asp440 and Leu438, undergo a rotamer change upon bromide binding. A second bromide peak, Br2, modeled at 48% occupancy, is located deep within the interdomain cavity. This ion makes contact with Ser318, Asn334, and Trp320, all contributed by the C-terminal domain. While residues surrounding both bromides are generally conserved in closely related CylK homologs, those associated with Br2 are not conserved in the C–O bond forming family member BrtB (Fig. S6C). This observation suggests that while Br2 residues might be relevant for paracyclophane forming enzymes, they may not be necessary for all family members.

### Key catalytic residues are located at Br1 within the active site cleft

Having discovered that the CylK active site cleft has two potential alkyl halide binding/activating regions, we next individually mutagenized several residues spanning these two locations, some of which were closely associated or appeared to directly bind either bromide. To interrogate specific roles for the residues associated with Br1 or Br2 in the native, paracyclophane forming reaction (Fig. 4C), we examined the ability of select variants to perform the first and second alkylation reactions with native substrates **1** and **2**. Considering that the first alkylation is effectively faster than the second (5), likely due to substrate and/or product inhibition, we monitored the first alkylation by interrogating the 1 hour time point of the native reaction, when wild-type enzyme has fully transformed both equivalents of **1** to intermediate **2** (>99% conversion). Notably, substitution of the residues that comprise the Br1 binding site (Arg105, Tyr473) abolished CylK activity towards **1** at 1 hour, while mutations that disrupt Br2 (Ser318, Asn334) had no apparent effect (Fig. 4D), retaining wild-type levels of activity. Variants targeting three additional residues located between the two bromide sites (Glu374, Leu438, Asp440) had very low or no activity and stood out as potential candidates for substrate binding or nucleophile activation due to their side chain functionalities and vicinity to Br1 on the cleft surface. Mutating Phe499 to Ala did not significantly reduce conversion.

We then examined the second alkylation reaction using a mixture of intermediate **2** containing a small amount of cylindrocyclophane F (∼18%) as the substrate. Consistent with the results obtained for the first alkylation reaction, variants that disrupted Br1 did not have appreciable activity on intermediate **2**, while substitution of residues comprising Br2 maintained near wild-type activity (Fig. 4E). Mutating Glu374, Leu438, and Asp440 also resulted in no significant activity toward **2** after 22 hours, while the Phe499 to Ala mutant retained appreciable activity as before. Of note, mutating Arg105 to Lys, and Asp440 to Asn resulted in no activity for either alkylation, suggesting that the size, hydrogen bonding capacity, and/or pK_a_ of these residues might be important for catalysis (Fig. S7). However, mutating Tyr473 to Phe only moderately reduced activity and might suggest that the aromatic nature and/or size of this residue is critical for alkylation. Based on these results, we propose that Br1 is the site of alkyl chloride binding and both aromatic ring alkylation events. Additional work is required to determine if the residues that comprise Br2 assist in substrate binding, but it is clear from these data that they are not essential for catalysis. Furthermore, the site of catalysis at Br1 is located near the outermost portion of the solvent exposed interdomain cavity; this is in agreement with previous work that demonstrated CylK’s ability to accept non-native substrates with rigid substitutions on the resorcinol ring nucleophile (8), which would necessitate a flexible or open active site.

### Molecular dynamics simulations suggest mode of substrate binding and activation

Having identified the site of alkylation and potential key residues, we sought to determine the specific mode of substrate binding and to assess the relative contributions of specific side chains towards nucleophile (resorcinol) versus electrophile (alkyl chloride) activation. We began by computationally docking two equivalents of native substrate **1** into the active site of CylK (Fig. 5A). The initial complexes were generated manually using the Br1 site to anchor the electrophilic alkyl chloride substituent of one equivalent of **1**. Previous DFT calculations performed on a model alkyl resorcinol substrate suggested interaction between a resorcinol phenol substituent and a carboxylate hydrogen bond acceptor would enhance nucleophilicity of the aromatic ring (8). Accordingly, potential interactions with essential carboxylate residues Asp440 and Glu374 guided the placement of the nucleophilic resorcinol of the other equivalent of substrate **1**. Additionally, we oriented the nucleophilic and electrophilic carbon centers at an appropriate distance (<4 Å) and arrangement to accommodate the known stereochemical inversion of the substitution reaction. Two docked initial complexes were used, and both were run as restrained and unrestrained simulations. Restrained simulations included 1 kcal/mol·Å^2^ restraints on all protein atoms to allow substrates to search the active site. All simulations consistently demonstrated that the active site cleft can accommodate two equivalents of substrate **1**, and furthermore, Asp440 and Glu374 maintained H-bonding contacts with the resorcinol nucleophile (Fig. 5B). For the majority of the simulation time, the chloride binding pocket formed by Arg105 and Tyr473 (Br1) maintained contact with the alkyl chloride electrophile (Fig. S8-S9). We conclude that Asp440 and Glu374 are likely resorcinol nucleophile activating residues, while Arg105 and Tyr473 bind and activate the alkyl chloride electrophile with hydrogen bonding interactions. Furthermore, the source of CylK’s exquisite regio- and stereoselectivity can be explained by the orientation in which the resorcinol nucleophile is held in close proximity to the backside of the carbon-chlorine (C–Cl) bond. Based on these results we can propose a mechanism of alkylation that is distinct from chemical and enzymatic precedent (Fig. 5C). Simulations of intermediate **2**, positioned for the second alkylation reaction, maintained similar alkyl chloride electrophile interactions, however, the resorcinol nucleophile did not remain at an appropriate distance or orientation for the reaction (Fig. S10). More work is required to accurately model the second alkylation step, however, it is clear from our mutagenic work that the same residues are essential for both alkylation events.

**Figure 5.**
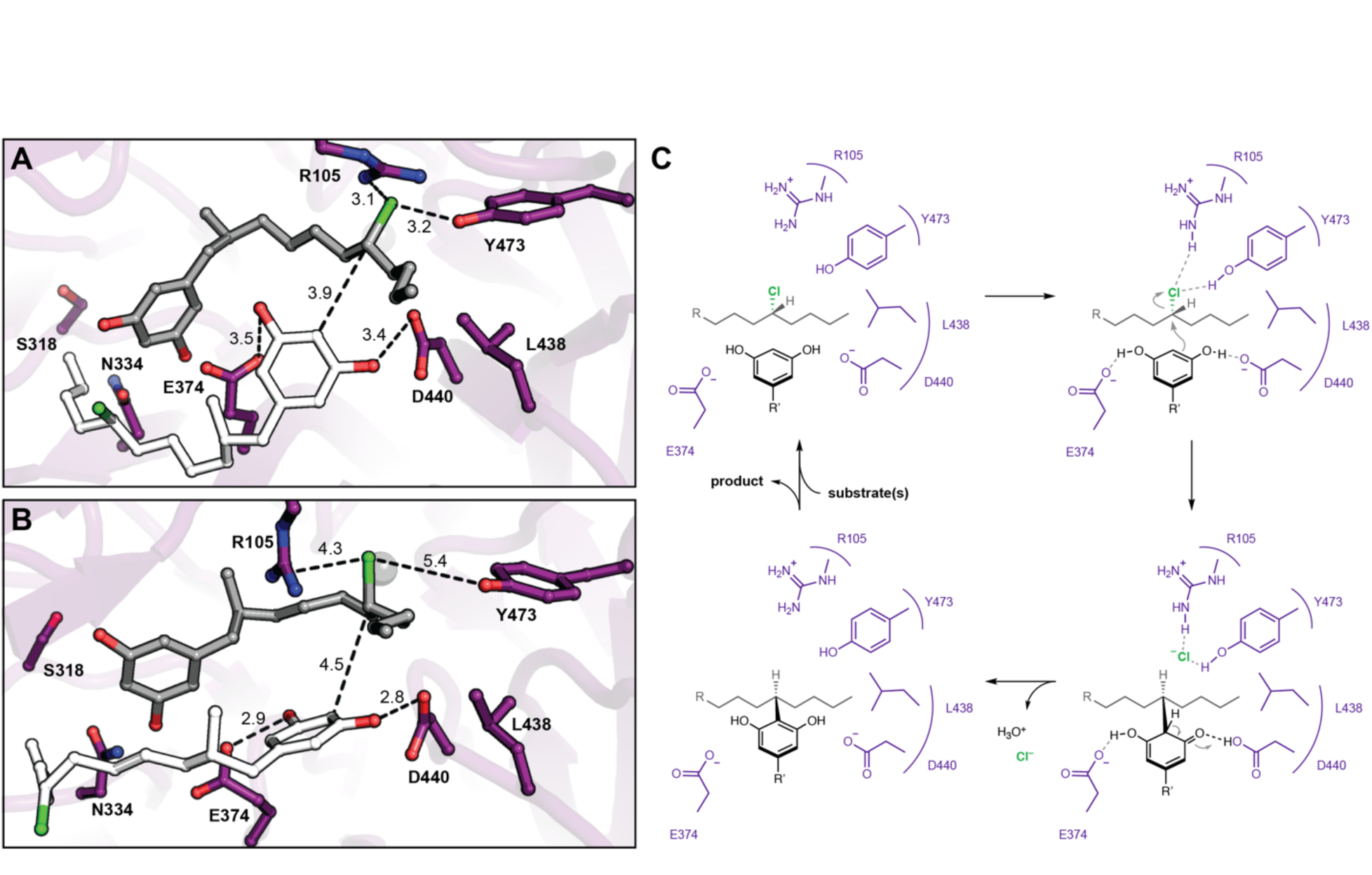
Molecular dynamics simulations reveal roles for key catalytic residues and enable a mechanistic proposal. (*A*) An energy minimized docking model of CylK in complex with two chlorinated alkylresorcinol molecules in cavity 1 before and (*B*) after unrestrained molecular dynamics simulation. This analysis shows that both substrates can be accommodated while maintaining contact with one another and essential catalytic residues. Substrate molecules and selected amino acid side chains shown in stick format. (*C*) Proposed mechanism for a single cycle of the CylK-catalyzed Friedel–Crafts alkylation with key residues illustrated.

### Bioinformatic analysis supports proposed residues implicated in alkyl chloride activation

We next sought to apply the functional insights gained from our structural analysis of CylK to improve our understanding of less closely related family members. Previously, bioinformatically identifying CylK-like enzymes was challenging because proteins encoding β-propeller motifs are common in publicly available protein databases, making it unclear which hits were true family members. With the knowledge that both protein domains of CylK are necessary for catalysis, we could now accurately locate CylK-like enzymes encoded in sequenced genomes. From a BLAST search, we identified over 700 proteins with >24% amino acid identity to CylK or BrtB (Dataset S1). That group was pared down to 286 unique enzyme sequences containing both N- and C-terminal domains found in CylK, and their relationship was assessed by constructing a maximum-likelihood phylogenetic tree (Fig. 6).

**Figure 6.**
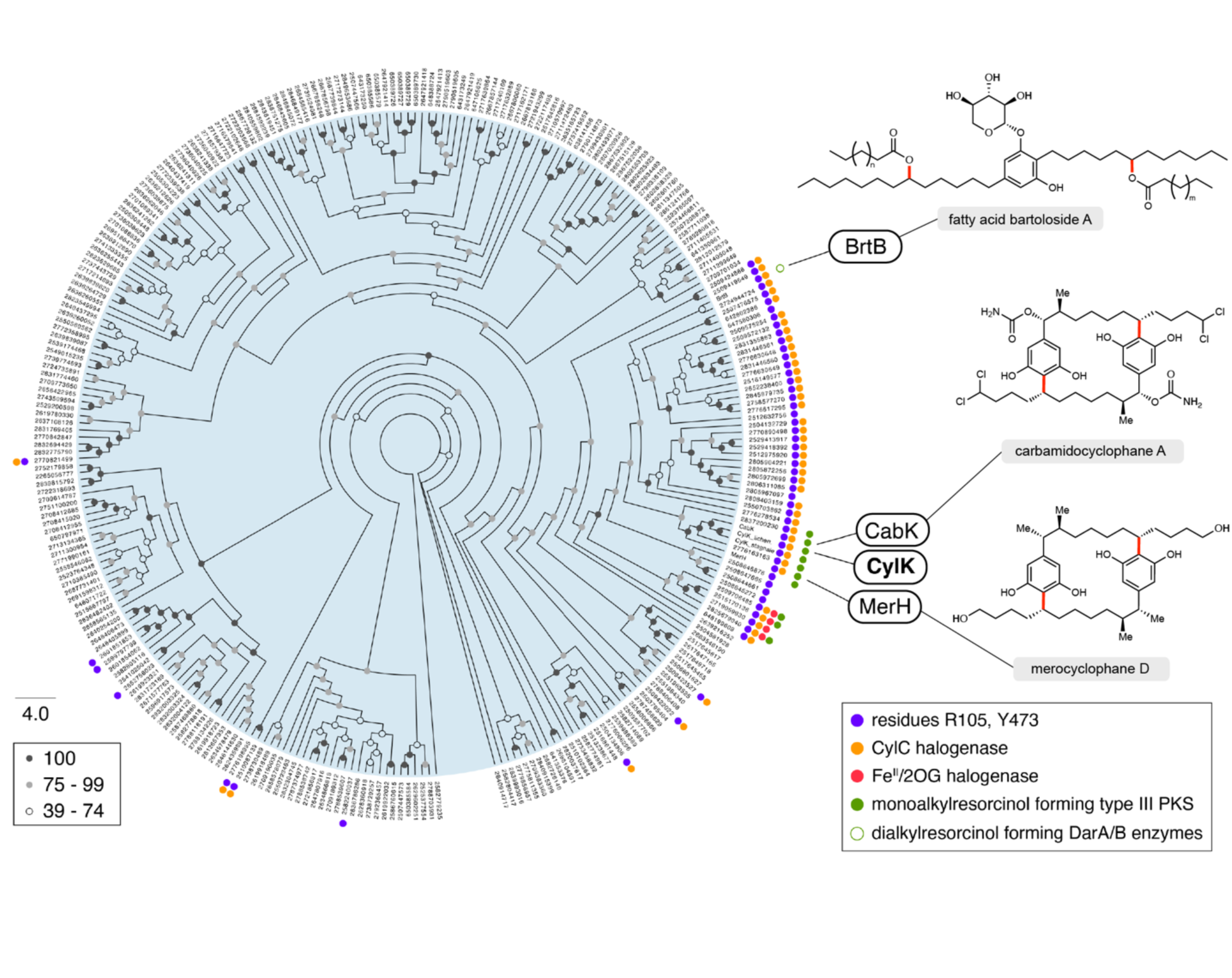
CylK homologs with partner CylC halogenases contain proposed alkyl chloride activating residues. Maximum-likelihood phylogenetic tree of protein sequences homologous to CylK and BrtB that contain both N- and C-terminal domains (>24% amino acid identity), highlighting key conserved catalytic residues, clustered halogenases, and associated monoalkylresorcinol forming enzymes. Dialkylresorcinol forming enzymes (DarA/B-like) were not found except as expected with BrtB and the bartoloside producing cluster, indicated as an unfilled green circle. Bootstrap values are shaded based on the legend. Natural products associated with select CylK homologs are displayed; the bonds highlighted in red are constructed by their respective CylK.

We further analyzed these data by looking for the presence or absence of a partner CylC halogenase encoded in the same genome. A subset of CylK homologs which are mostly clustered together on the phylogenetic tree are found in organisms that also have a putative CylC halogenase. Of these CylK and CylC enzyme pairs, ∼80% are co-localized in their respective genomes, hinting that they might work together within a biosynthetic pathway (Dataset S2). We found very few other types of halogenases co- localized with CylK homologs; those identified were of the iron(II) 2-(oxo)-glutarate (Fe^II^/2OG)-dependent enzyme family but were also clustered with CylC halogenases. We examined each putative CylK sequence for the presence of the proposed alkyl chloride activating residues Arg105 and Tyr473. Intriguingly, nearly all of the candidate CylK enzymes containing both key residues were from organisms that also encode a CylC halogenase (42 of 53 organisms), suggesting that they are likely using halogenated substrates and that Arg105 and Tyr473 are indeed important for alkyl chloride activation. This subset of CylK homologs was aligned and visualized (Fig. S11) which revealed that, while the resorcinol nucleophile-activating residue Asp440 was also highly conserved, Leu438, Glu374, and other active site residues are not highly prevalent within this subset. This demonstrates that the correlation between Arg105, Tyr473, and the presence of a CylC halogenase is not simply due to the overall similarity of this subset of enzymes.

Inspired by the biosynthetic logic of the *cyl* pathway, we also looked for co-localized monoalkylresorcinol (MAR) forming enzymes encoded near putative CylK homologs in order to ascertain if resorcinols are likely to serve as their native nucleophilic substrates. To our knowledge, biosyntheses of MARs only involves type III PKS enzymes such as CylI. (24) Surprisingly, we found only 10 CylI homologs clustered with CylK homologs, representing just 19% of the candidate enzymes that have Arg105 and Tyr473 residues and are expected to have alkyl chloride activating activity. This observation suggests that a broader diversity of nucleophilic substrates might be used by uncharacterized alkyl chloride activating enzymes. Of note, dialkylresorcinol forming enzymes such as those involved in bartoloside biosynthesis were not found in CylK-encoding gene clusters beyond the bartoloside gene cluster. The observation that the majority of CylK homologs do not have proposed alkyl chloride activating residues, partner halogenases, nor resorcinol forming enzymes, indicates that this enzyme family may catalyze other reactions not involving alkyl halides or resorcinols. This information will enable future studies by prioritizing organisms that might produce more distantly related natural products and/or CylK-like enzymes with alternate substrate scopes.

## Discussion

Our results provide a structural basis for understanding the unusual enzymatic Friedel–Crafts alkylation catalyzed by CylK in cylindrocyclophane biosynthesis. This structural model, together with supporting biochemical experiments and bioinformatic analyses, have provided fundamental insight into this intriguing reaction that represents the only known example of aromatic ring alkylation with an alkyl halide electrophile. The molecular knowledge we have gained may be applied to access and engineer novel biocatalysts, design chemocatalysts, and discover new natural products that may be candidate therapeutics and/or play important physiological roles in cyanobacteria.

We identified the active site of CylK at the interface of its N- and C-terminal domains. Together with previously characterized enzymes, our results show how structurally related β-propeller enzymes have divergently adapted this fold for catalytic function. In two metalloenzymes, carotenoid cleavage dioxygenase (CCD) (25) and nitrous oxide reductase (NOR) (26), an active site metallocofactor is lodged on the top face of the propeller and coordinated by His side chains from inner β-strands of the fold (Fig. S4). Like CylK, both of these enzymes have a capping motif on the top face of the propeller. CCD contains more extensive internal loops than CylK, and these adopt structured helix-loop motifs that bury the top face of the propeller. NOR is a di-domain dimeric enzyme that employs an electron transfer domain from an adjacent monomer to shield the catalytic metallocofactor in the β-propeller motif. In both, these domain arrangements create a favorable environment for substrate binding and catalysis, as we observe in CylK (Fig. S12).

Both domains of CylK have several bound calcium ions, a row of which are near the proposed alkyl chloride binding site. However, the closest Ca^2+^ ion is approximately 8 Å from this pocket. Although we have previously demonstrated a functional requirement for calcium (5), our structural work suggests that a direct role in catalysis is unlikely. Removing calcium might cause a significant conformational change that precludes access to or alters the architecture of the active site, as the treatment of CylK with EDTA (ethylenediaminetetraacetic acid) was observed to cause a shift in oligomeric state from monomer to a dimer. This might be indicative of larger structural changes within each unit of CylK that are incompatible with catalysis. It is unknown if calcium plays a regulatory role *in vivo* although it is a component of the cyanobacterial BG-11 growth medium.

We propose a plausible mechanism for CylK catalysis that invokes positioning the reactive partners in close proximity for a concerted S_N_2-like reaction, consistent with the known inversion of configuration at the alkyl chloride stereocenter upon substitution (Fig. 5C). In particular, we suggest that the side chain carboxylates of Asp440 and Glu374 hydrogen bond with the two phenol substituents of the resorcinol nucleophile. By deprotonating one phenol, the essential residue Asp440 could enhance the nucleophilicity of the reactive carbon. Analogous deprotonation of a resorcinol has been invoked as a critical step in an enzymatic Friedel–Crafts acylation reaction with acetyl arene electrophiles, lending support to our proposal (27). Simultaneously, we hypothesize that Arg105, Tyr473, Phe499, and the hydrophobic portion of Thr84 form an alkyl chloride binding pocket as observed in the bromide soaked CylK structure. Hydrogen bonding interactions between the polar residues and the alkyl chloride could weaken the C–Cl bond, reducing the overall activation energy barrier for substitution. Interestingly, mutating Tyr473 to Phe only slightly reduces alkylation activity which suggests that the alkyl chloride Tyr473 interaction might resemble an anion-pi interaction (28, 29). While there is precedent for alkyl halide-arene interactions (30), they are less thoroughly studied than anion-pi interactions. Alternatively, if Tyr473 provides a key hydrogen bond, the Phe mutant may partially compensate by forming an alkyl halide-pi interaction. In either case, this alkyl chloride binding pocket is reminiscent of the active sites of the *S*-adenosylmethionine (SAM)-dependent chlorinase (31) and fluorinase (32) enzymes. In these structures, similar ensembles of polar and aliphatic residues bind and stabilize a halide anion as it reacts with SAM in a S_N_2 substitution reaction that resembles the reverse of the transformation catalyzed by CylK (Fig. S13A-B). Interestingly, our proposed CylK alkyl chloride binding mode is distinct from that of the well-studied haloalkane dehalogenase enzyme, which primarily uses the indole N–H bonds of adjacent Trp residues to activate a primary alkyl chloride for alkylation by the side chain carboxylate of an active site Asp via hydrogen bonding (Fig. S13C). (33) We speculate that the differences between these two enzymes arise from their distinct evolutionary histories as well as the decreased hydrophobicity and increased reactivity of a primary alkyl chloride substrate. Following alkylation, the chloride anion product diffuses out of the active site, and the aromaticity of the resorcinol is restored upon proton transfer to water, perhaps via His 391. Finally, intermediate **2** is released from the active site, must reorient 180°, and re-bind for the second intramolecular alkylation to occur in the same fashion. Although we cannot rule out a radical mechanism, the architecture of the active site and lack of radical initiation sources suggests that one-electron processes are unlikely.

We have emphasized the distinction between the proposed resorcinol nucleophile-activating residues and those that likely activate the alkyl chloride electrophile in order to compare CylK with the divergent C–O bond forming family member, BrtB (30% amino acid identity, 46% similarity). We hypothesize that both enzymes utilize a similar strategy to enhance the reactivity of their alkyl chloride substrates, although it remains to be determined if BrtB is similarly stereoinvertive. In agreement with this proposal, BrtB contains Arg105 and Tyr473 equivalents that may play a role in alkyl chloride activation. In place of the nucleophile- activating Asp440, BrtB contains a functionally similar Glu, and notably, Glu374 is substituted with a positively charged Arg. These two residues might interact with carboxylate nucleophiles, although the precise roles of these residues in BrtB’s distinct reactivity remain to be determined. Interestingly, a carboxylate nucleophile would be predominantly deprotonated at physiological pH, suggesting that nucleophile activation might be unnecessary in this enzyme. Regardless, the divergent reactivity of CylK and BrtB hints that other relatives might perform unique chemistry and may even be amenable to protein engineering efforts to alter their substrate scopes and reactivity.

The reaction catalyzed by CylK represents a unique approach for enzymatic C–C bond formation. Other biological strategies to form analogous arene-alkyl C–C bonds involve radical intermediates that must be precisely held and oriented in their enzyme active sites in order to prevent side reactions. Examples include the radical SAM and Fe^II^/2OG enzymes involved in streptide (34) and etoposide (35) biosynthesis, respectively. In contrast to CylK, these enzymes form bonds between arenes and unactivated carbon centers. Because the target sp^3^ coupling sites of their substrates are inherently unactivated, these alkylating enzymes must stringently discriminate between the alkyl C–H coupling site and its chemically equivalent neighbors. In contrast, the CylK catalyzed reaction leverages the preinstalled alkyl chloride substituent in order to direct reactivity. Therefore, the cylindrocyclophane biosynthetic logic can be described as dividing the task of determining reaction specificity between CylK and its partner halogenase, CylC. This novel halogenase family likely has strict requirements for substrate positioning (36), although it remains to be seen if CylC is promiscuous. This feature of distributing selectivity and alkylation across two enzymes might rationalize why wild-type CylK can accept multiple non-native substrates, whereas active site specificity for unactivated coupling partners might be more stringent.

CylK performs a challenging chemical reaction without precedent in biology. By elucidating its structure, we have begun to unravel the molecular details underlying the use of alkyl halides as cryptic intermediates in natural product biosynthesis. This enzymatic Friedel–Crafts alkylation might also find application in constructing diverse alkyl-arene C–C bonds outside of natural pathways. With structural information about CylK, notably the identification of active site residues, efforts to further expand its substrate specificity and improve its stability via enzyme engineering can begin in earnest. In addition to its unique reactivity, the large size and solvent accessibility of CylK’s active site suggest it might be engineered to accept diverse and bulky substrates. Furthermore, our structure and proposed mechanism have increased our fundamental understanding of enzymatic interactions with alkyl halides and may inspire the design of biomimetic chemocatalysts that engage secondary alkyl halides in substitution reactions (37, 38). Finally, armed with the ability to functionally annotate CylK-like genes, we can prioritize orphan biosynthetic gene clusters and producing organisms in order to discover additional structurally unique natural products.

## Materials and Methods

### Protein expression and purification

CylK was expressed and purified for crystallographic characterization according to a modified literature method (8). Following overexpression in *E. coli* BL21(DE3) as a SUMO fusion construct, cell pellets were resuspended in lysis buffer (20 mM Tris pH 8.0, 500 mM NaCl, 10 mM MgCl_2_, 10 mM CaCl_2_) supplemented with an EDTA-free Pierce Protease Inhibitor Tablet. Cells were lysed using a continuous-flow homogenizer (Avestin EmulsiFlex-C3). The pellet was collected by centrifugation (20,000*g* for 30 min) and resuspended in lysis buffer supplemented with 6 M urea and 5 mM imidazole. The denatured enzyme solution was clarified twice by centrifugation (20,000*g* for 15 min) and the supernatant was applied to 20 mL of Ni NTA resin. The column was washed with lysis buffer containing 6 M urea and 5 mM imidazole. Bound species were eluted with lysis buffer containing 6 M urea and 250 mM imidazole. All subsequent steps were carried out at 4 °C. The enzyme was refolded by sequentially dialyzing this solution against buffer 1 (lysis buffer supplemented with 3 M urea) overnight, buffer 2 (lysis buffer lacking urea) for 4 h, and buffer 3 (lysis buffer supplemented with 5% glycerol) for 4 h. Following dialysis, the refolded enzyme solution was centrifuged (100,000*g* for 30 min) and the supernatant was collected and incubated overnight with ULP Protease (1:25 w/w). Then, following a subtractive Ni NTA step, the enzyme was applied to a Superdex 200 30/100 gel filtration column (GE Healthcare) equilibrated in protein storage buffer (20 mM HEPES pH 7.8, 50 mM NaCl, 10 mM MgCl_2_, 10 mM CaCl_2_, 10% glycerol). The fractions containing pure CylK were pooled, concentrated (500-700 µM, determined by extinction coefficient and absorbance at 260 nm) and flash frozen in liquid nitrogen.

### Crystallization and structure determination

Prior to crystallization trials, purified CylK protein in storage buffer was combined with 5-undecylbenzene-1,3-diol **3**, a substrate analog synthesized as described previously (8) and dissolved in protein storage buffer containing 22% DMSO. The final protein solution contained 10 mg/mL CylK and 1.66 mM of **3**. Crystals were obtained by using the hanging drop vapor diffusion method in 2 μL drops mixed in 1:1 ratio with a precipitant solution of 1.8 M sodium malonate, pH 7.0. To aid in obtaining phase information, the native Ca(II) sites in CylK crystals were substituted with a lanthanide via soak in a solution of 2.0 M sodium malonate, 100 mM terbium(III) chloride for 8 min at room temperature. The Tb-soaked crystals were transferred to a cryoprotectant solution containing 2.0 M sodium malonate supplemented with 20% ethylene glycol, mounted on rayon loops, and flash frozen by direct plunge into liquid nitrogen. Diffraction datasets were collected at 1.6314 Å (Tb anomalous) and 0. 97872 Å (native) on these crystals at beamlines 23-ID-B [General Medical Sciences and Cancer Institutes Collaborative Access Team (GM/CA-CAT), Advanced Photon Source (APS)] and 21-ID-F [Life Sciences Collaborative Access Team (LS-CAT), APS], respectively. Diffraction datasets were processed with the HKL2000 software package (39). The structure was solved by using the MR-SAD method. PHASER-EP (40, 41), as implemented within the PHENIX software package (42), was used to calculate initial phases. A poly-alanine model generated from the core β-propeller domain of *Staphylococcus cohnii* streptogramin B lyase (PDB accession code 2QC5, with all loops and secondary structure connections truncated) (19) was used as the initial search model for molecular replacement. PHENIX.autobuild generated an initial model containing 504 residues in 15 chain fragments with initial R_work_/R_free_ values of 29.9%/37.4%. The model was iteratively improved via refinement in Refmac5 against a native dataset and model building in Coot (43), yielding R_work_/R_free_ values of 16.4%/19.8% (Table S1). The final model contains residues 7-45, 49-392, and 413-662 with 11 Ca^2+^ ions, 2 Mg^2+^ ions, 1 Cl^-^ ion, and 339 water molecules. Tb ions were not modeled because they were not present at high occupancy. Although **3** was required for CylK crystallization, and weak electron density resembling the compound could be found at several points near the surface of the protein, likely mediating crystal lattice contacts, we could not confidently model the alkyl resorcinol. No density for **3** could be identified in cavity 1, the putative active site of CylK.

To identify possible alkyl chloride-binding sites within the protein, CylK crystals were soaked in 2 M sodium malonate with 500 mM NaBr for up to 1 min. The Br-soaked crystals were transferred to a cryoprotectant solution consisting of the soak solution supplemented with 20% ethylene glycol. Diffraction datasets were collected at 0.9184 Å (Br anomalous) and 0.97872 Å (native) on these crystals at beamlines 23-ID-B (GM/CA-CAT, APS) and 21-ID-F (LS-CAT, APS), respectively. Phase information was obtained by molecular replacement using the CylK model as the initial search model in Phaser MR, implemented within CCP4. The model was iteratively built and refined in Coot and Refmac5, respectively, resulting in a R_work_/R_free_ of 16.9%/19.8% (Table S4). The final model contains residues 7-393 and 410-667 with 10 Ca^2+^ ions, 2 Mg^2+^ ions, 1 Na^+^ ion, 4 Br^−^ ions, and 403 water molecules. Anomalous maps were generated using CAD and FFT programs implemented within CCP4. Figures were generated in PyMOL (Schrödinger, LLC).

### Mutant enzyme activity screening

CylK mutants were expressed and assayed for activity according to a modified literature method (8). Point mutants in plasmid pPR-IBA1-CylK were synthesized and sequence verified by GENEWIZ (South Plainfield, NJ, USA), DNA sequences available (Fig. S14). Enzyme activity assays using substrate pair **3** + **4** were carried out as follows. Chloro-SNAc electrophile **4** was accessed as described previously (8). Respective mutant, wild-type, and negative control plasmids (Addgene #31122), were freshly transformed into electrocompetent *E. coli* BL21 Gold CodonPlus® (DE3) RIL cells (Agilent), and an individual colony was used to directly inoculate 1 mL cultures of LB medium supplemented with 100 μg/mL ampicillin or carbenicillin and 34 μg/mL chloramphenicol in a 96 deep well plate (VWR), sterility was maintained with standard techniques and a gas permeable rayon film (VWR). The liquid culture plate was incubated for 5 hours at 37 °C and 250 rpm shaking. After five hours, and confirming visible growth in each well, the liquid culture plate was cooled to 15 °C, maintaining shaking. Following an additional 30-45 minutes of incubation, protein expression was induced with 250 μM IPTG. The cultures were incubated for 4 h at 15 °C with 250 rpm shaking. Following expression, 10 μL of each culture was removed from each well and sub-cultured for liquid culture RCA-based DNA sequencing to confirm identity and rule out culture cross contamination. The remaining cultures were concentrated ∼10X by centrifugation (3,220*g* for 10 min) and resuspended in ∼200 μL of spent media supernatant. Reactions were initiated in opaque 96-well plates (Costar) by combining 100 μL of concentrated cell suspension with 2 μL of lysozyme mix (9 mg lysozyme & 1 mg EDTA-free Pierce Protease Inhibitor Tablet per 500 μL of 500 mM Tris, 50 mM EDTA, pH 8.0) and mono-alkylation substrates in DMSO for a final concentration of nucleophile **3** at 150 μM, electrophile **4** at 300 μM, and DMSO at 3.4%. Reaction plates were sealed with aluminum seals (VWR) and incubated at 37 °C with 190 rpm shaking for 20 hours. Reactions were quenched with the addition of 1:1 methanol/acetonitrile (200 μL) and centrifuged (3,220*g* for 15 min). The supernatant was then subjected to LC-MS (Liquid Chromatography-Mass Spectrometry) to measure ion abundance of product **5** (C_31_H_53_ClNO_4_S [M + H^+^] = 536.3768 ± 5 ppm) and starting material **4** (C_14_H_26_ClNO_2_S [M + H^+^] = 308.1446 ± 5 ppm). LC-MS conditions as published previously (8). Reactions were repeated in biological triplicate.

Enzyme activity on native substrates **1** or **2** was assayed as follows. Substrate **1** and intermediate **2** were accessed chemoenzymatically as described previously (5). Care was taken to minimize glycerol in substrate stocks because it was determined to inhibit CylK activity. Respective mutant and wild-type plasmids were freshly transformed into electrocompetent *E. coli* BL21 Gold CodonPlus® (DE3) RIL cells (Agilent), and 5-10 colonies were used to directly inoculate 100 mL cultures in LB medium supplemented with 100 μg/mL ampicillin or carbenicillin and 34 μg/mL chloramphenicol. The liquid cultures were incubated at 37 °C and 190 rpm shaking. At OD400 = 0.4, the cultures were cooled to 15 °C, maintaining shaking. Following an additional 1 hour, protein expression was induced with 250 μM IPTG. The cultures were then incubated for 4 h at 15 °C with 190 rpm shaking. Following expression, cell pellets were collected by centrifugation (3,220*g* for 10 min) and resuspended in ∼2 mL of assay buffer (20 mM HEPES pH 7.8, 100 mM NaCl, 10 mM MgCl_2_, 5 mM CaCl_2_). Cell concentration was normalized across mutants by measuring OD600 and diluting with an appropriate amount of assay buffer. A 600 μL aliquot of each mutant was lysed by sonication (Branson, 25% amplitude, 1 min) on ice. The soluble fraction was collected by centrifugation (16,100*g* for 20 min) and analyzed by anti-Strep-HRP (IBA) western blotting to confirm mutant solubility and approximate protein concentration (Fig. S15). Reactions were initiated by combining 4.3 μL of **1** or **2** with 25.7 μL of soluble lysate for a final concentration of **1** at 400 μM and 4% DMSO, and **2** at 200 μM and 2% DMSO. Reactions were sealed and incubated at 37 °C for 1 or 20 hours. Reactions were quenched with the addition of 1:1 methanol/acetonitrile (60 μL) and centrifuged (6,000*g* for 15 min). The supernatant was then subjected to analytical HPLC (High Performance Liquid Chromatography) to determine percent conversion as described previously. Reactions were repeated in biological duplicate, and then repeated again on a second day to minimize any potential variation. Results were consistent on both days.

### Molecular Dynamics Simulations

Coordinates for the native substrates **1** and **2** were prepared in ACEDRG (44) from MOL file descriptions of the molecules generated in ChemDraw. Two substrate molecules were docked manually into cavity 1 of CylK using Br1 to place the alkyl chloride component and interactions with essential amino acids to guide placement of the second substrate. The resulting CylK complexes were prepared for molecular dynamics (MD) simulations using the Xleap module of AmberTools (45). Substrates were parameterized using Antechamber (46, 47) in AmberTools, while the FF14SB forcefield (48) was used for protein atoms. Complexes were solvated in a 10 Å octahedral box of in TIP3P water (49). Sodium and chloride ions were added to neutralize the complexes and achieve a final concentration of 150 mM NaCl. All minimizations and production runs were performed with Amber20 (45). Minimization was performed using 5000 steps of steepest descent and 5000 steps of conjugate gradient. In the first round, 500 kcal/mol·Å^2^ restraints on all protein, ligand atoms as well as the metal ion cofactors. Restraints were reduced to 100 kcal/mol·Å^2^ for an additional round of solvent minimization. Next, restraints were retained only ligands and metal ions for a third round of minimization. Finally, all restraints were removed for the last stage of minimization. After minimization, a 100 ps run was used to heat complexes from 0 to 300 K using constant volume periodic boundaries with 10 kcal/mol·Å^2^ restraints on all solute atoms. Equilibration was performed using two 10 ns runs (10 kcal/mol·Å^2^ and 1 kcal/mol·Å^2^ restraints) on protein, ligand and metal ions. Following equilibration, simulations were run as either restrained or unrestrained. For restrained simulations, we obtained 100 ns trajectories with 1 kcal/mol·Å^2^ restraints on protein atoms. Unrestrained simulations (100 ns) were run with all restraints removed.

### Bioinformatic Analysis

The CylK (GenBank: ARU81125.1) and BrtB (GenBank: AOH72618.1) protein sequences were independently used in a Basic Local Alignment Search Tool (BLAST) search of the Joint Genome Institute-Integrated Microbial Genomes & Microbiomes (IMG-JGI) database of all isolates, metagenome-assembled genomes (MAGs), and single-amplified genomes (SAGs). Each BLAST search resulted in 500 hits (>24% amino acid sequence identity) and were merged to form a non-redundant list of 715 enzyme candidates (Dataset S1). Parent enzyme sequences (CylK, BrtB) and likely CylK homologs (MerH, CabK, CylK from *C. stagnale*) were added to this dataset. A targeted BLAST search of the genomes from this list was performed to identify all putative halogenases (CylC-like, Fe^II^/2OG, SAM-dependent, flavin-dependent, & haloperoxidase) and resorcinol forming enzymes (CylI, BrtC, BrtD). All non-CylC halogenases were confirmed to contain key catalytic residues and examined for co-localization with CylK. All candidate CylC halogenases were included, although the overwhelming majority (63 of 78 enzymes) were fewer than 18 genes away from CylK. (Dataset S2). CylC’s catalytic residues remain unknown. Monoalkylresorcinol forming CylI homologs were included if clustered with CylK; 11 of 54 candidate enzymes were less than 12 genes away from CylK (Dataset S2). Next, candidate CylK enzymes were filtered by constructing an individual MUSCLE alignment (50, 51) of each hit to the CylK parent sequence and identifying at least 135 amino acid residues before the C-terminal domain. Lys240 or its aligned equivalent was defined as the junction point between protein domains. This analysis resulted in 286 unique protein sequences (Dataset S2) that were aligned (MUSCLE, EMBL-EBI), sites containing gaps at ≥ 90% of sequences were removed, and a maximum-likelihood phylogenetic tree was constructed using IQ-TREE v1.6.12 (52) and visualized with FigTree v1.4.4. The best fit model (VT+F+G4) was selected by ModelFinder (53). Bootstrap values were calculated using the UFBoot2 method (54). These candidate enzymes were analyzed for the presence or absence of Arg105 and Tyr473 residues by constructing an individual MUSCLE alignment of each hit to the CylK parent sequence. The results were manually mapped onto the phylogenetic tree. The subset CylK homologs with both residues were MUSCLE aligned and visualized with ConSurf (55). Code used to perform iterative MUSCLE alignments available (https://github.com/nbraffman/CylK-homologs).

## Acknowledgments

We gratefully acknowledge use of the resources of the Advanced Photon Source, a US Department of Energy (DOE) Office of Science User Facility operated for the DOE Office of Science by the Argonne National Laboratory under Contract DE-AC02-06CH11357. Use of Life Sciences Collaborative Access Team Sector 21 was supported by the Michigan Economic Development Corporation and the Michigan Technology Tri-Corridor (Grant 085P1000817). The National Institute of General Medical Sciences and National Cancer Institute Structural Biology Facility at the Advanced Photon Source (GM/CA@APS) has been funded in whole or in part with federal funds from the National Cancer Institute (ACB-12002) and the National Institute of General Medical Sciences (AGM-12006, P30GM138396). The Eiger 16M detector at GM/CA@APS was funded by NIH Grant S10 OD012289. We thank Andrew J. Mitchell and Jonathan A. Bergman for their excellent technical assistance. This work was supported by NSF Grants 1454007 and 2003436, the Cottrell Scholar Award (to E.P.B.), and NIH Grant GM119707 (to A.K.B.).

## SI Appendix

**Figure S1.**
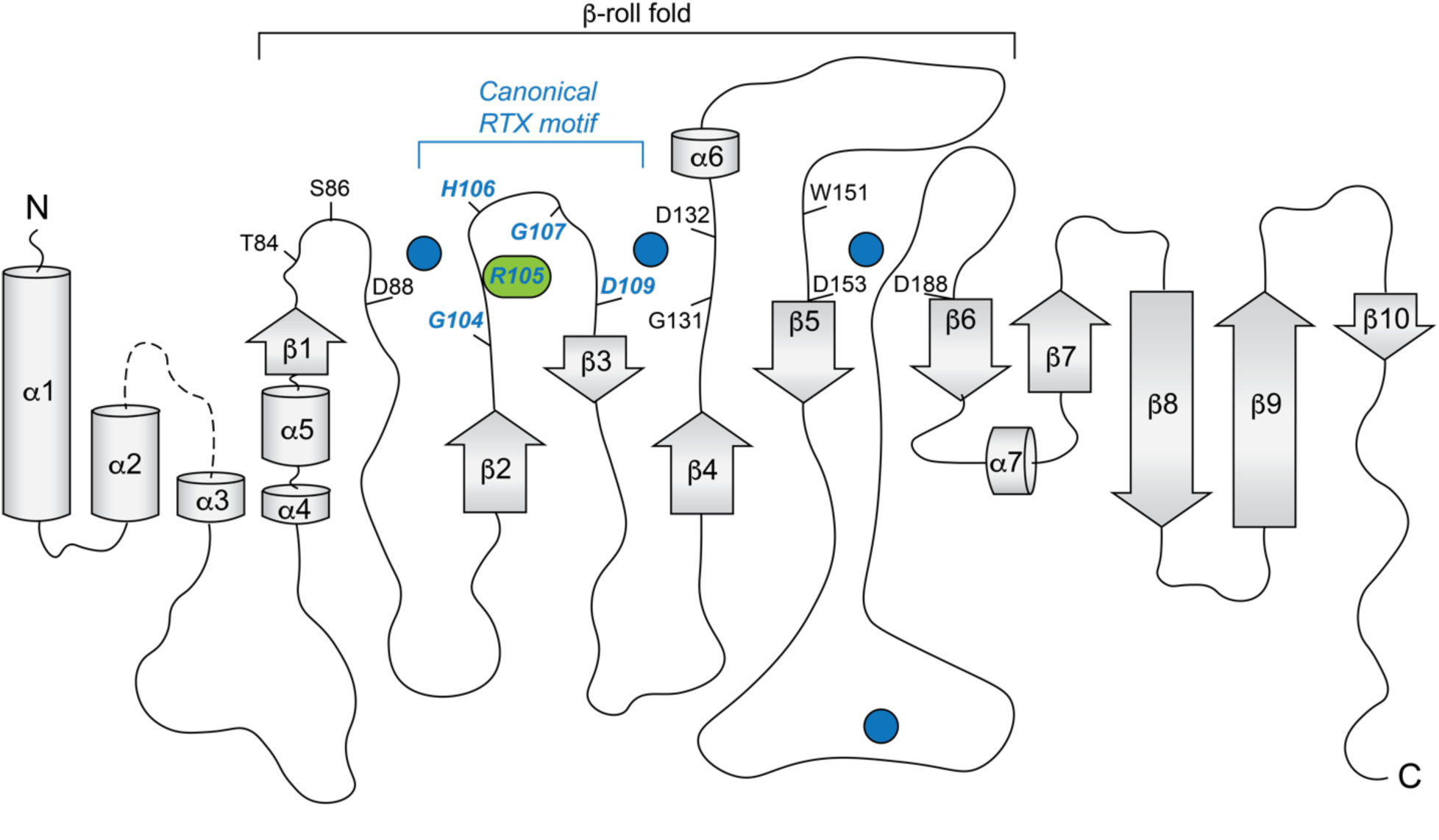
Topology diagram of the N-terminal CylK β-roll domain, residues 7-251. Ca^2+^ ions are shown as blue circles, residues implicated in catalysis are highlighted in green ovals, Ca^2+^-coordinating residues are shown in black, and Ca^2+^-coordinating residues that follow canonical repeat-in-toxin (RTX) motifs are shown in blue italics.

**Figure S2.**
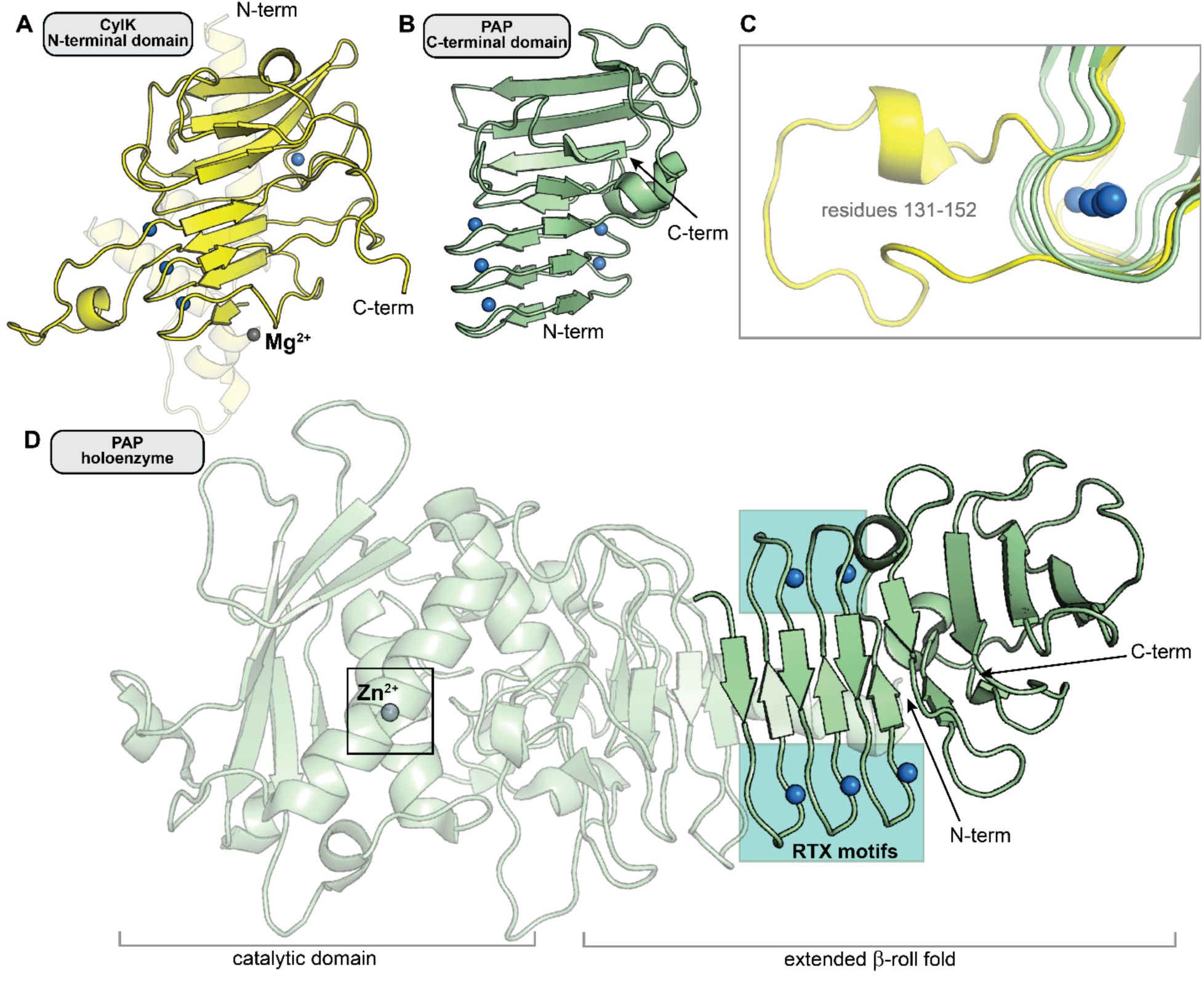
Comparison of the N-terminal CylK β-roll domain to similar C-terminal repeat-in-toxin (RTX) domains in other enzymes. *(A)* The N-terminal domain of CylK (residues 7-251). *(B)* The C-terminal region of RTX-motif enzyme psychrophilic alkaline metalloprotease (PAP) that structurally aligns with the β-roll of CylK (residues 326-461) (PDB ID 1OMJ). *(C)* Some CylK Ca^2+^-coordinating loops (yellow, residues 131-152) extend beyond the tight RTX-motif turn (green, PAP residues 330-335, 348-353, and 366-371). *(D)* The Ca^2+^-binding β-roll motif in PAP is located far away from the active site of the enzyme (box) and is likely not involved in catalysis (1). In all panels, Ca^2+^ ions are represented as blue spheres. Mg^2+^ (CylK) and Zn^2+^ (PAP) ions as shown as grey spheres.

**Figure S3.**
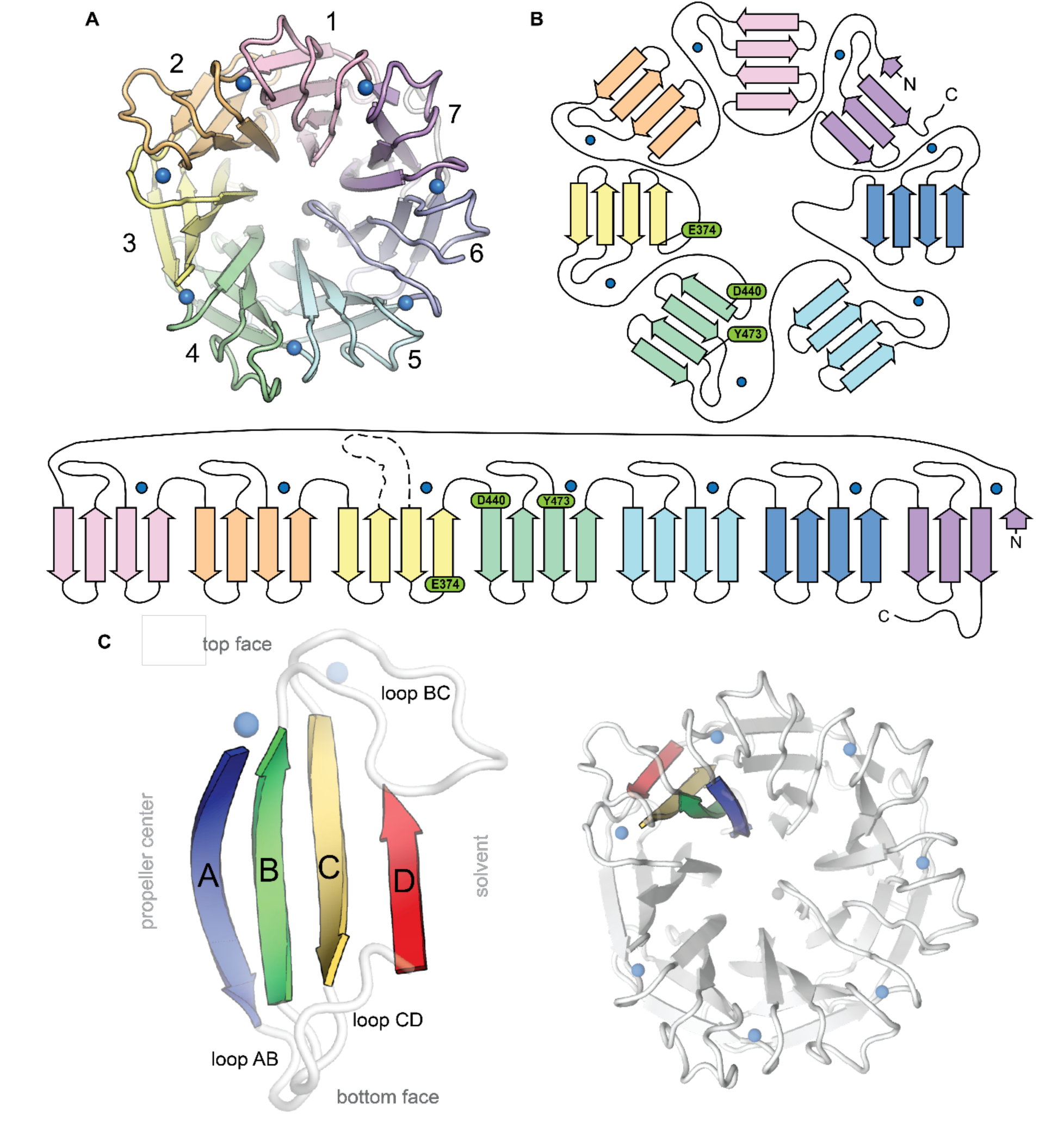
Topological and structural features of the C-terminal CylK β-propeller domain. *(A)* Top-down view of the C-terminal β-propeller domain (residues 252-662). *(B)* Topology diagrams of the CylK β- propeller domain. Blades are colored as in (A), Ca^2+^ ions shown as blue circles. Residues implicated in catalysis are shown in green. *(C)* Example of a propeller blade (left) and the orientation of the blade within the β-propeller fold (right). β-strands within each blade are lettered A-D, with A corresponding to the innermost strand, closest to the central tunnel. Loops are labeled to indicate the strands they connect. For example, loop BC connects blade B and C within the blade while loop DA connects strand D of one blade to strand A of the next blade.

**Figure S4.**
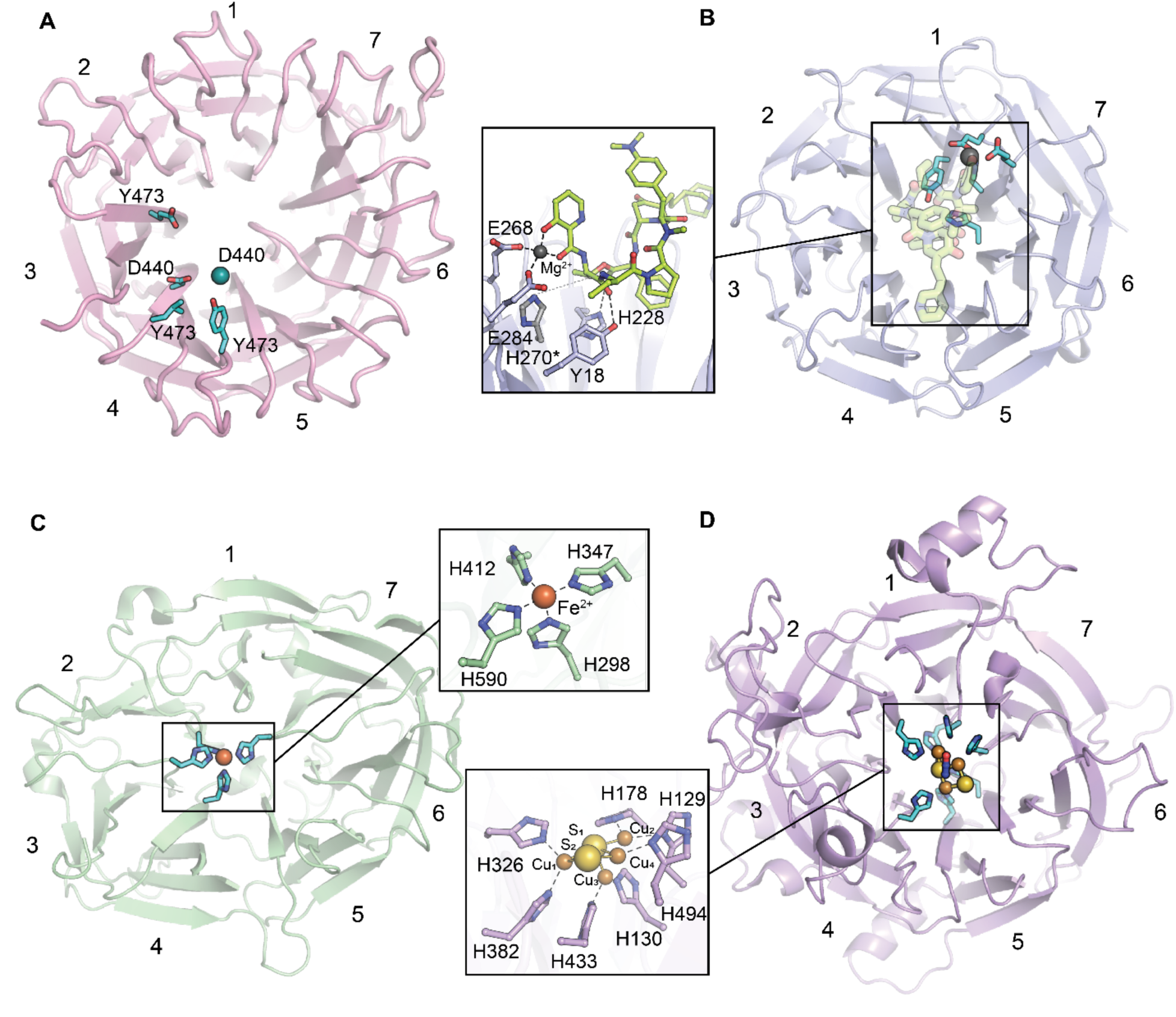
Comparison of the proposed CylK active site location to those of other characterized 7-bladed β-propeller enzymes. Panels show a top-down ribbon diagram of β-propeller domains. Residues implicated in catalysis are shown as cyan sticks. *(A)* The C-terminal domain of CylK. Br^-^ ions are shown as teal spheres and Ca^2+^ ions are shown as transparent blue spheres. *(B)* Virginiamycin B lyase (VrgB) complexed with the substrate analog, quinupristin (lime green sticks) (PDB ID 2Z2P). Mg^2+^ ions are shown as grey spheres. *H270 modeled as in PDB ID 2Z2O. *(C)* Carotenoid cleavage dioxygenase (CCD) (PDB ID 3NPE). Fe^2+^ ions are shown as orange spheres. *(D)* Nitrous oxide reductase (NOR) complexed with N_2_O (blue and red sticks) (PDB ID 3SBR) and the active site Cu-S cluster. Cu ions are shown orange spheres, and sulfur atoms are shown as yellow spheres.

**Figure S5.**
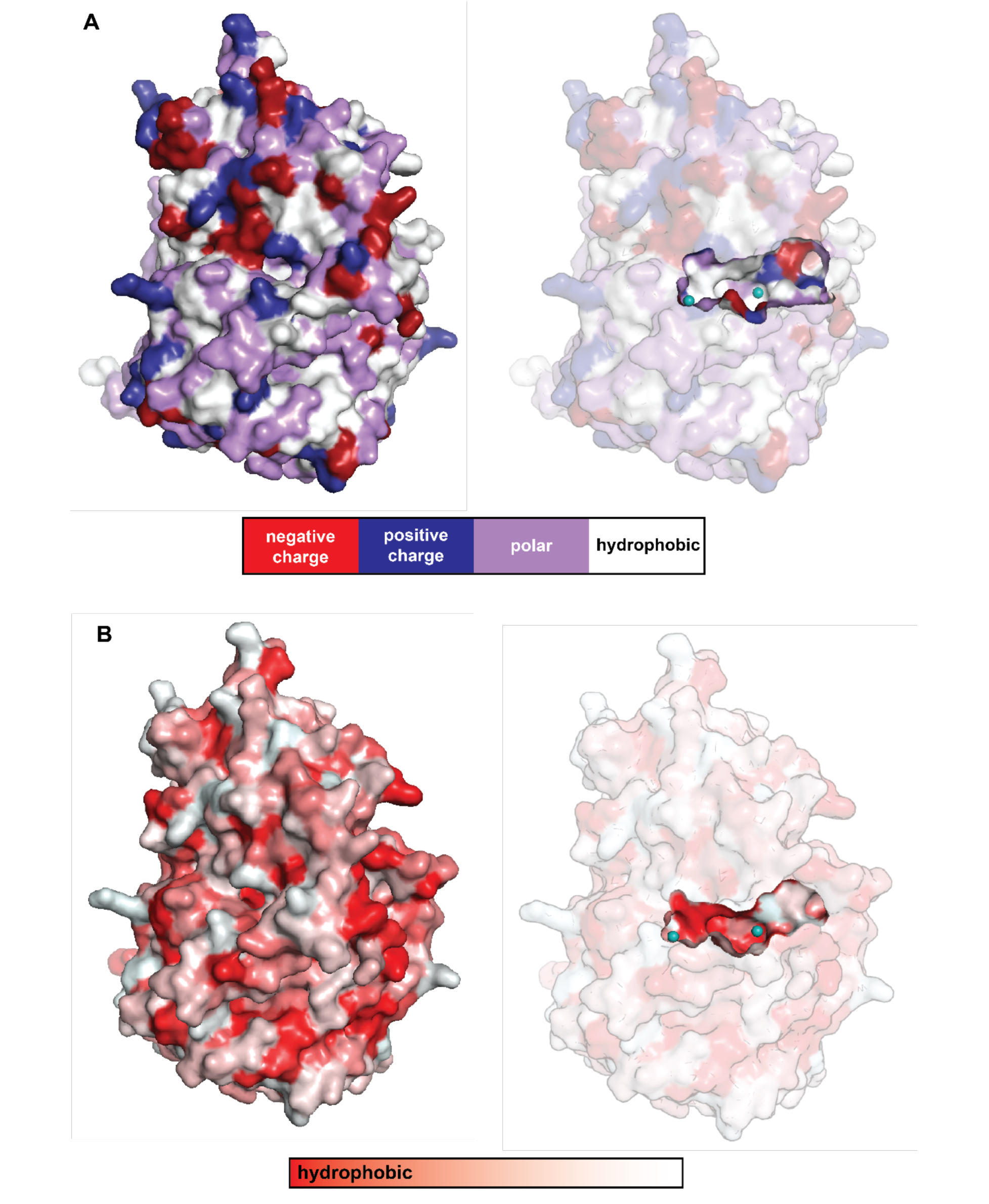
Solvent accessibility and properties of the cavity located in between the C-terminal β-propeller domain and the N-terminal domain in CylK. The walls of this cavity contain *(A)* charged (red/blue) and polar (purple) residues and *(B)* hydrophobic patches (red), ideal for interaction with the amphipathic resorcinol.

**Figure S6.**
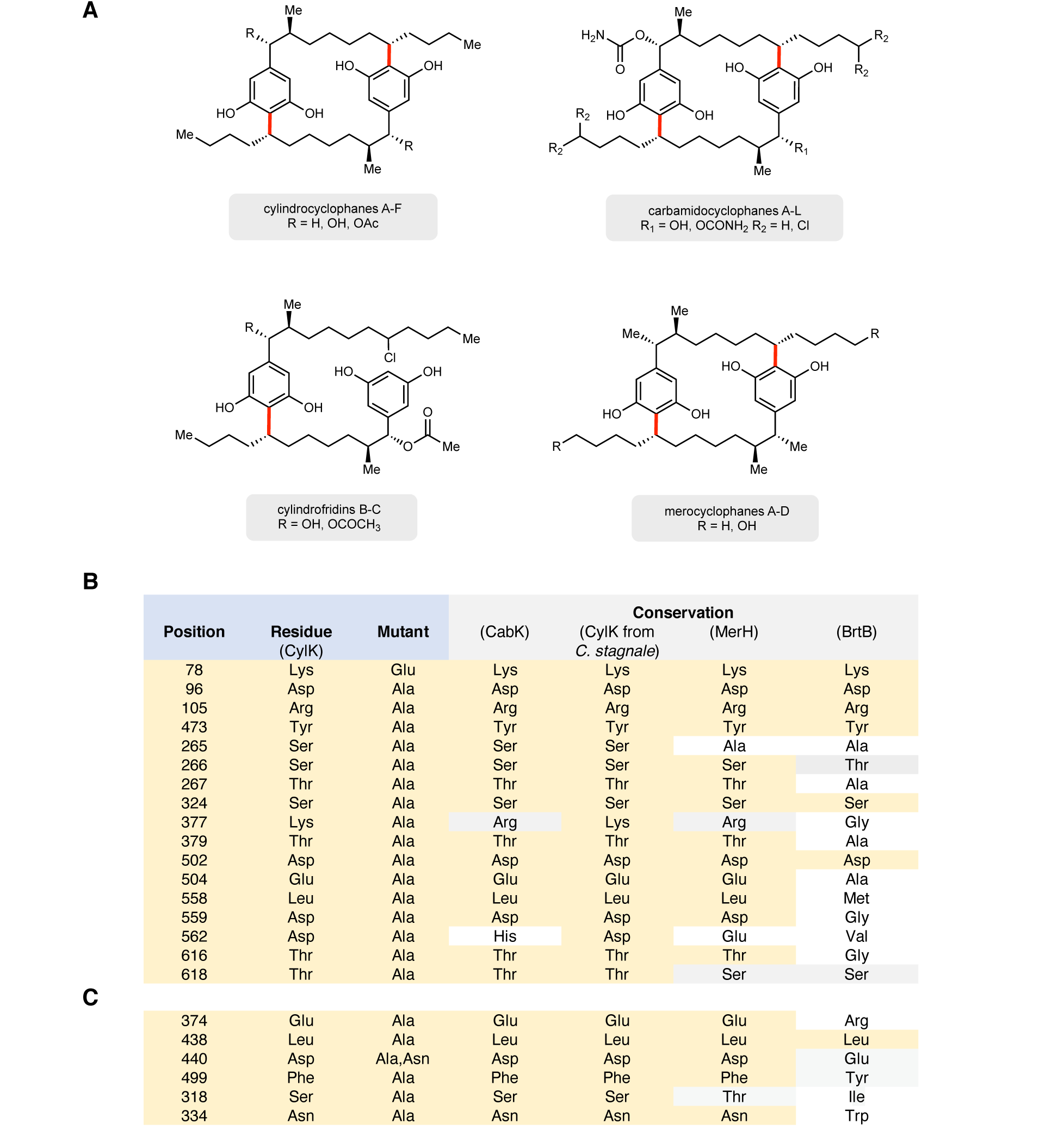
Cylindrocyclophane-like natural products and their corresponding CylK homologs guide mutagenesis. *(A)* Structurally related metabolites produced by cyanobacteria with related biosynthetic gene clusters and CylK homologs. Namely, merocyclophanes (*Nostoc* sp. UIC 10110, MerH), cylindrofridins (*Cylindrospermum stagnale* PCC 7417, CylK), and carbamidocyclophanes (*Nostoc* sp. CAVN2, CabK). *(B)* Conservation of CylK residues selected for initial mutagenesis. Polar, conserved residues of CylK were mutated to alanine or glutamate in order to biochemically determine the location of the active site. *(C)* Conservation of CylK residues within the active site cleft selected for second round of mutagenesis.

**Figure S7.**
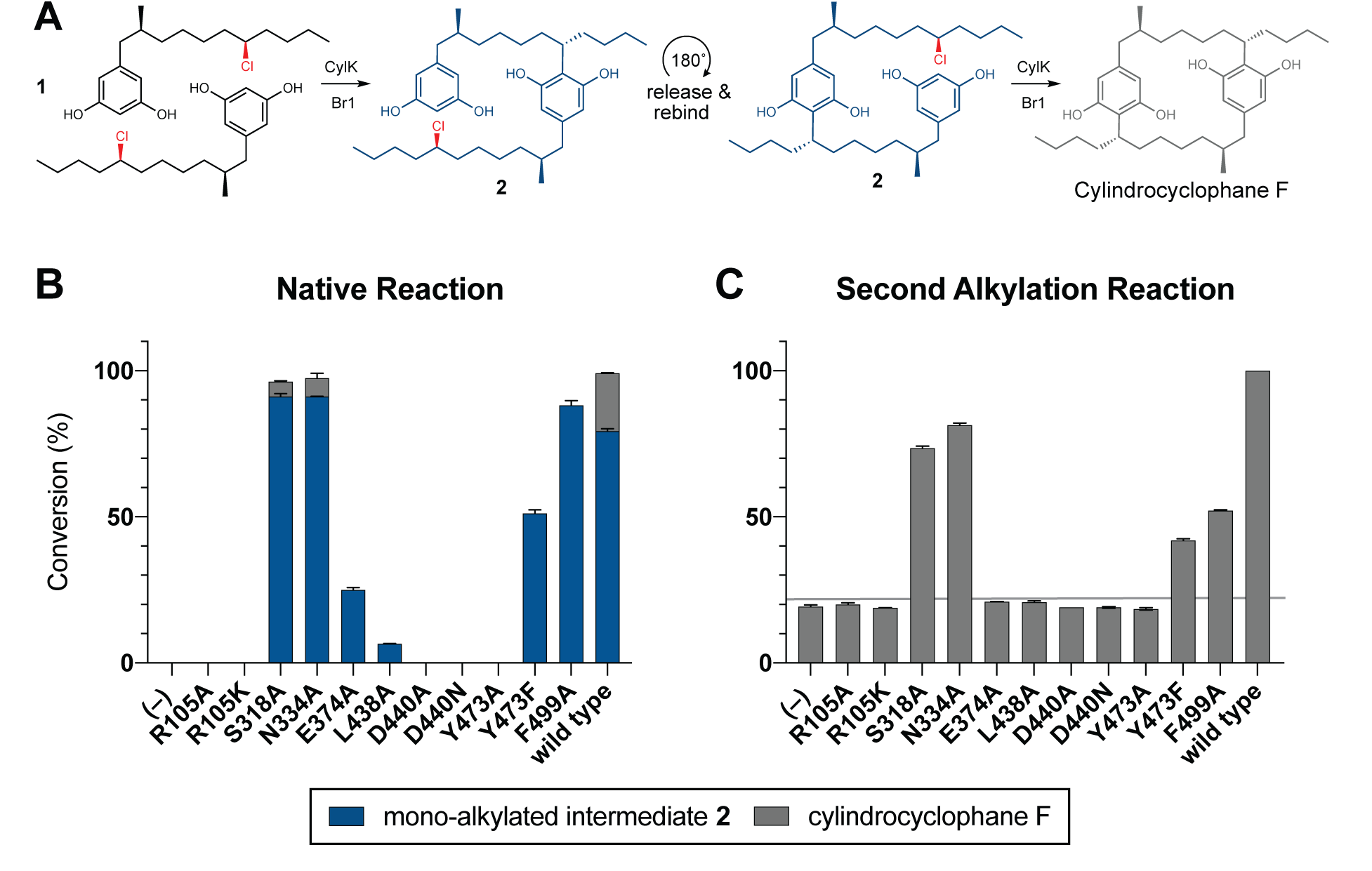
Mutagenesis of active site cleft. (*A*) Proposed alkylation scheme invoking release and rebinding of mono-alkylated intermediate **2** for the second alkylation event towards cylindrocyclophane F. (*B*) End- point activity at 1 hour of select mutants performing the native reaction with substrate **1**. Product formation was quantified by HPLC, error bars represent the standard deviation from the mean of two biological replicates. (*C*) End-point activity at 22 hours of select mutants performing the second alkylation reaction with intermediate **2**.

**Figure S8.**
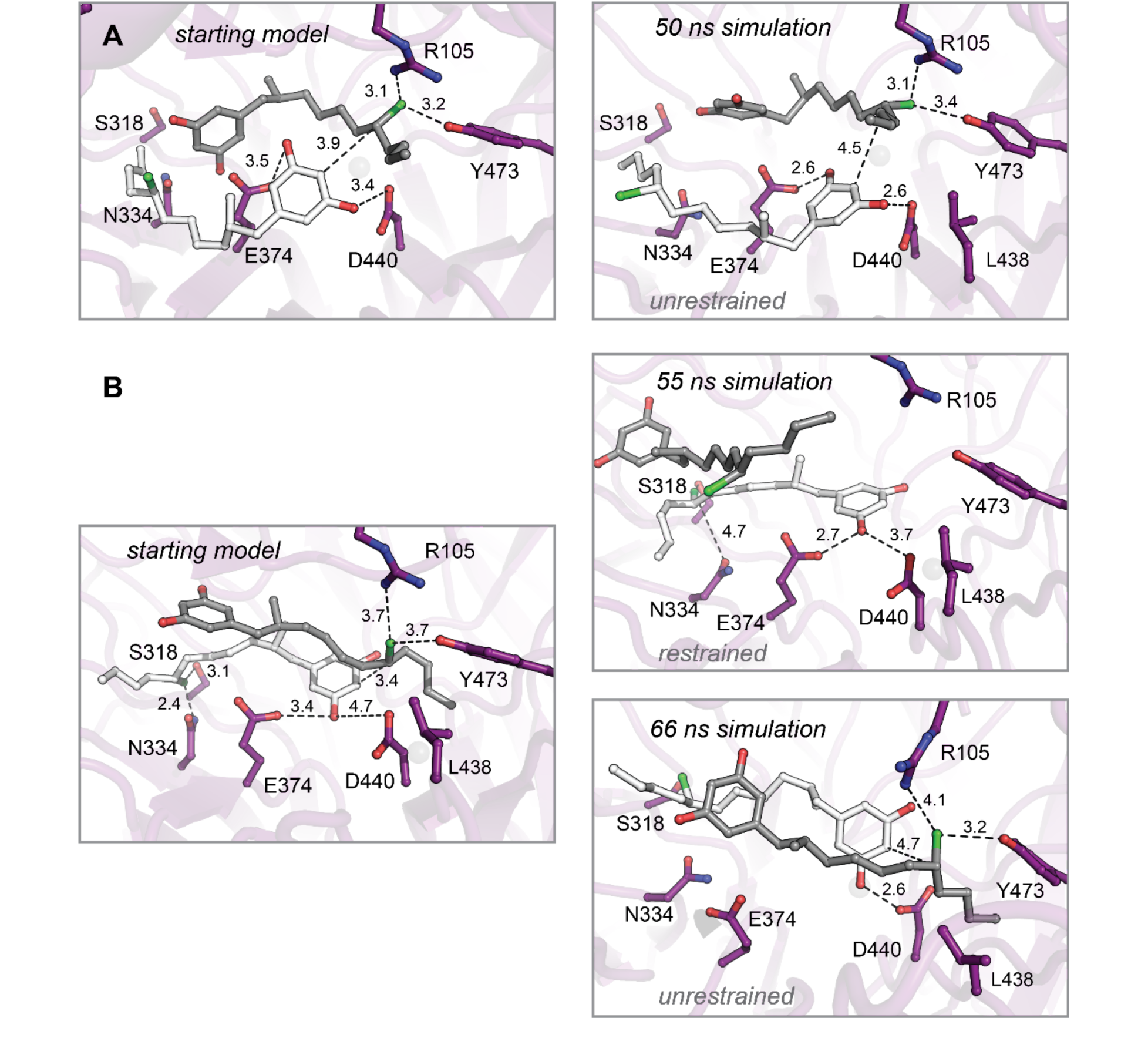
Molecular dynamics simulations of enzyme-substrate complexes manually docked into interdomain cavity support active site assignment in this region of CylK. *(A)* An energy minimized starting model before (left) and after restrained molecular dynamics simulation (right). The complex was generated using Br1 to place one equivalent of substrate **1** (gray). This complex also maximizes interaction between the resorcinol of a second substrate molecule (white) and amino acids D440 and E374 in the propeller domain. Results from an unrestrained simulation are shown in the main text (Fig. 4D-E). *(B)* Restrained (top right) and unrestrained (bottom right) simulations resulting from a starting model (left) that prioritized Br2 site interactions. Select residues shown as dark purple sticks, distances shown as dashed lines and measured in angstroms.

**Figure S9.**
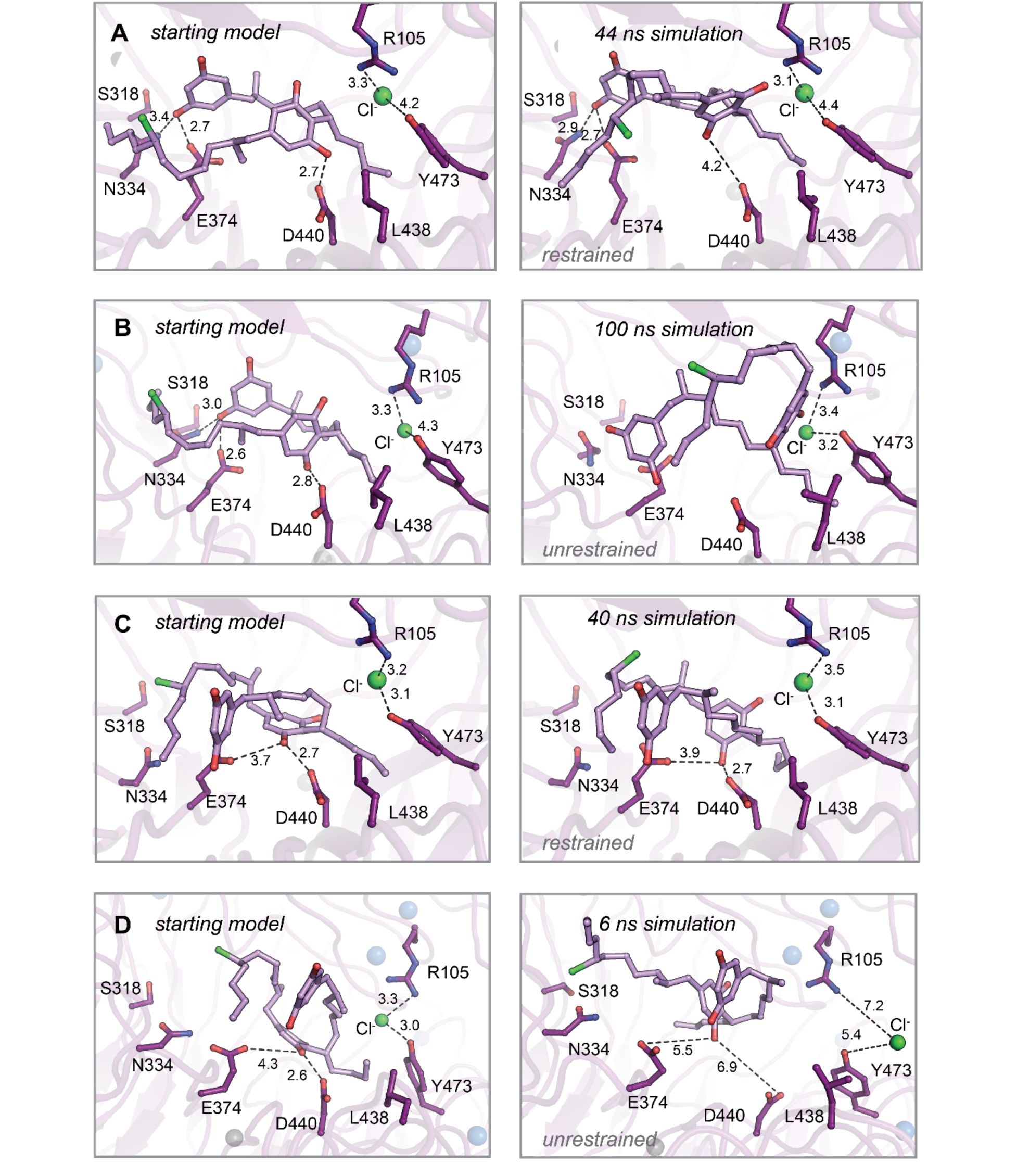
Molecular dynamics simulations of intermediate product **2** complexes manually docked into interdomain cavity reveal a halide binding pocket involving R105 and Y473. Energy minimized starting models before (left) and after simulation (right). *(A)* Restrained and (*B*) unrestrained simulations of the intermediate product complex. *(C)* Restrained and (*D*) unrestrained simulations of the intermediate product complex in an alternate starting conformation. Select residues shown as dark purple sticks, product intermediate shown as light purple sticks, distances shown as dashed lines and measured in angstroms.

**Figure S10.**
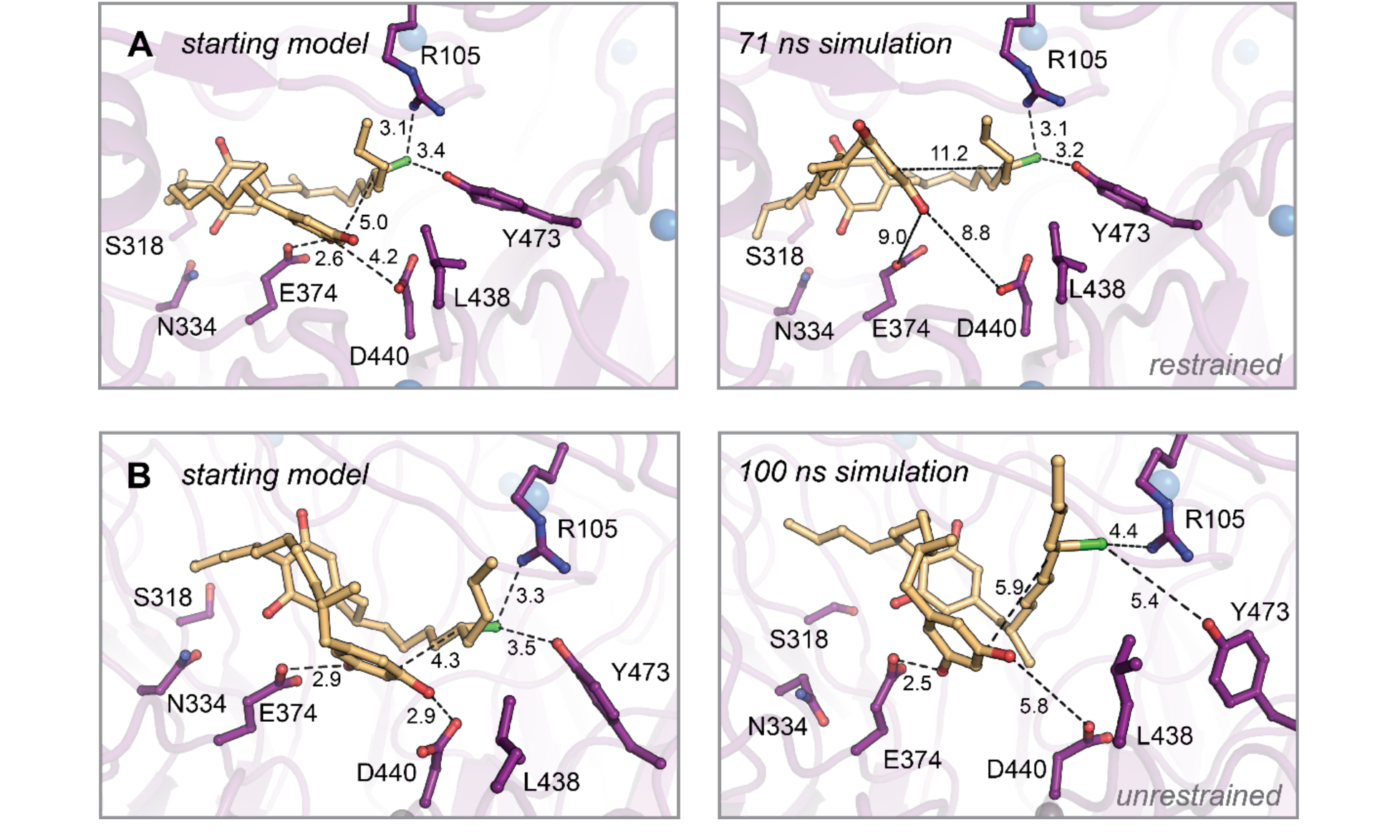
Molecular dynamics simulations of monoalkylated intermediate **2** positioned for the second alkylation reaction. Energy minimized starting models before (left) and after simulation (right). *(A)* Restrained and *(B)* unrestrained simulations of A. Select residues shown as dark purple sticks, monoalkylated intermediate **2** shown as orange sticks, distances shown as dashed lines and measured in angstroms.

**Figure S11.**
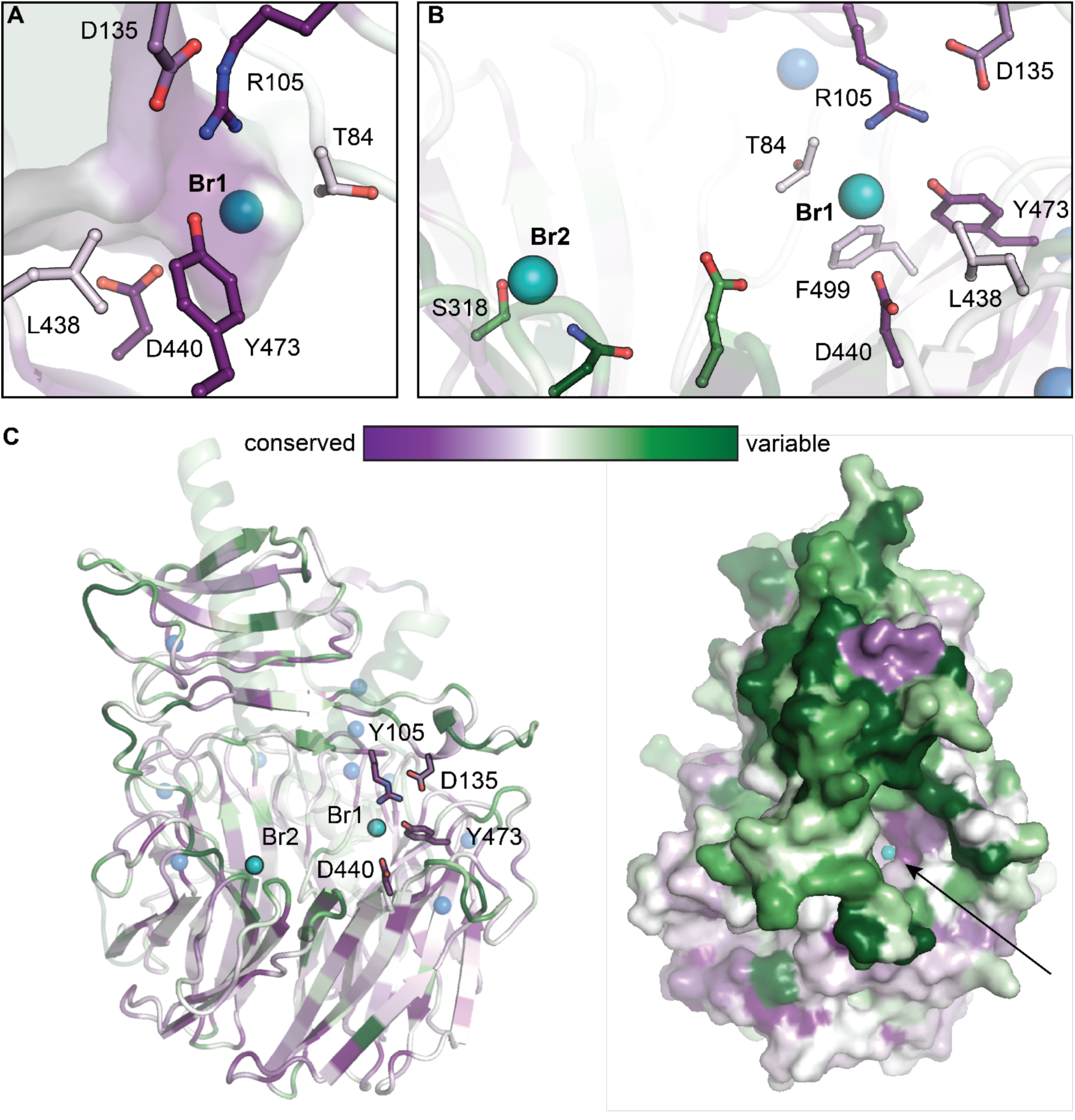
Conservation mapping analysis of functionally annotated CylK homologs. Representations of sequence conservation generated in ConSurf (2) using a subset of cyanobacterial CylK homologs that have R105 and Y473 (48 unique sequences). Of those homologous enzymes, 37 contain a partner CylC halogenase in the genome of the encoding organism (∼80% are co-localized with CylK). *(A)* The halide binding pocket in CylK homologs with R105 and Y473. *(B)* Residues surrounding Br1 are generally conserved in this subset, while Leu 438, Glu 374, and other active site residues are not highly conserved, potentially suggesting a diversity of substrates used by these candidate enzymes. Residues surrounding Br2 are variable. *(C)* Ribbon and surface diagrams of overall sequence conservation in this subset of homologs. Selected amino acid residues are represented in stick format, bromide ions are shown as teal spheres, and calcium ions are shown as blue spheres. Arrow denotes the opening of the interdomain cavity proposed to contain the active site of CylK.

**Figure S12.**
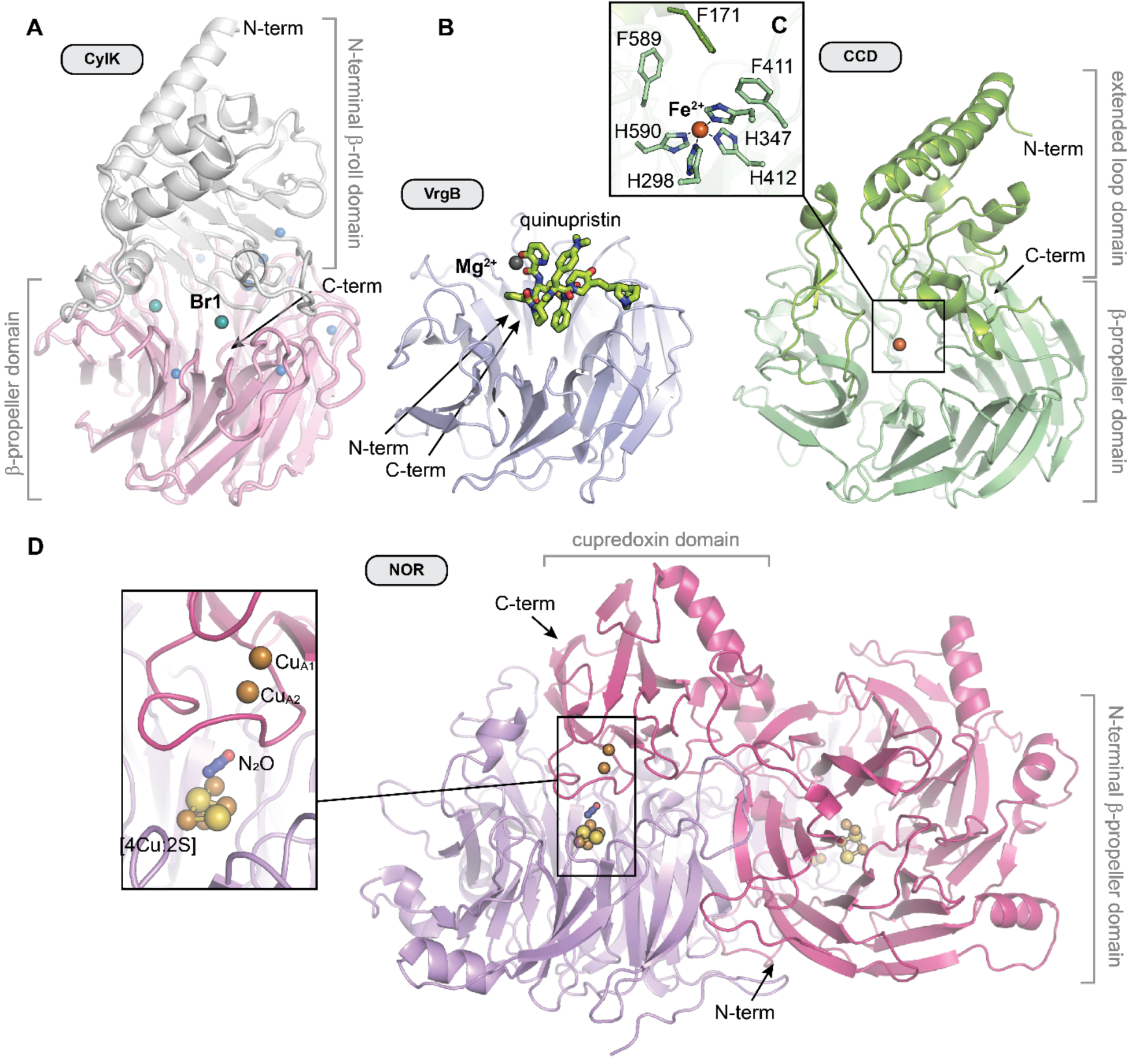
A comparison of the overall structures of CylK and selected 7-bladed β-propeller enzymes. *(A)* CylK is a fusion of a Ca^2+^-binding β-roll fold and a β-propeller with the active site located at the domain interface. *(B)* Virginiamycin B lyase (VrgB) is a single domain enzyme in which the active site is located on the top face of the propeller. Quinupristin substrate analog shown as green sticks (PDB ID 2Z2P). *(C)* The extended loops of carotenoid cleavage dioxygenase (CCD) form a helical domain on the top face of the propeller, oriented similarly to the N-terminal domain of CylK. The active site, delineated by the location of the iron cofactor (orange sphere), is located at the interface of these two domains (PDB ID 3NPE). *(D)* Dimeric nitrous oxide reductase (NOR) contains an active site at the interface between a C-terminal cupredoxin electron transfer domain (Cu_A_) contributed by the other monomer and the β-propeller catalytic domain (inset). Adapted from (3) (PDB ID 3SBR). Unless otherwise noted, ions are shown as spheres and colored as in Fig. S2.

**Figure S13.**
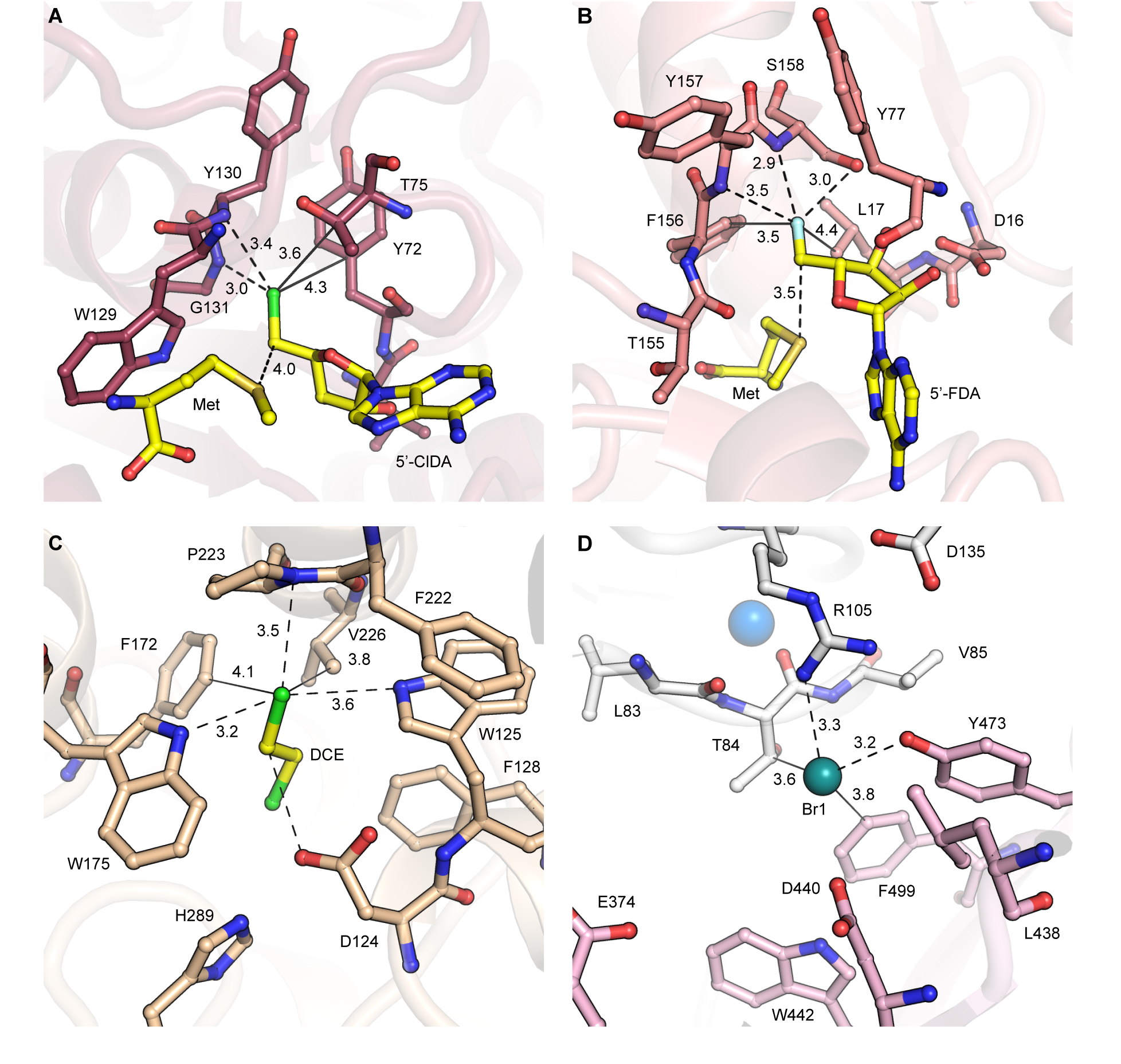
Comparison of bound alkyl halides and their respective enzyme active sites. *(A)* 5’-chloro-5’- deoxyadenosine (5’-ClDA) and methionine (Met) products bound in the active site of wild-type SalL SAM-dependent chlorinase (PDB ID 2Q6I). The alkyl chloride forms apparent polar interactions (dashed lines) with the backbone amides of Gly 131 and Tyr 130; and non-polar interactions (solid lines) with Tyr 72 and Thr 75. *(B)* 5’-fluoro-5’-deoxyadenosine (5’-FDA) and methionine (Met) products bound in the active site of wild-type FlA SAM-dependent fluorinase (PDB ID 1RQR). The alkyl fluoride forms apparent polar interactions with the backbone amides of Ser 158 and Tyr 157, and the side chain of Ser 158; and non-polar interactions with Phe 156 and Leu 17. *(C)* 1,2-dichloroethane (DCE) substrate bound in the active side of wild-type haloalkane dehalogenase (PDB ID 2DHC). The alkyl chloride forms apparent polar interactions with Trp 125, 175, and the backbone amide of Pro 223; and non-polar interactions with Phe 172 and Val 226. Asp 124 is the nucleophilic residue that displaces chloride. *(D)* Bromide ion product analog bound in the active site of CylK. The Br^-^ ion forms polar interactions with Arg 105 and Tyr 473; and non-polar interactions with Thr 84 and Phe 499. Select amino acid residues are represented in stick format, bromide ions are shown as teal spheres, calcium ions are shown as blue spheres, alkyl chloride in green, and alkyl fluoride in light aqua.

**Figure S14.**
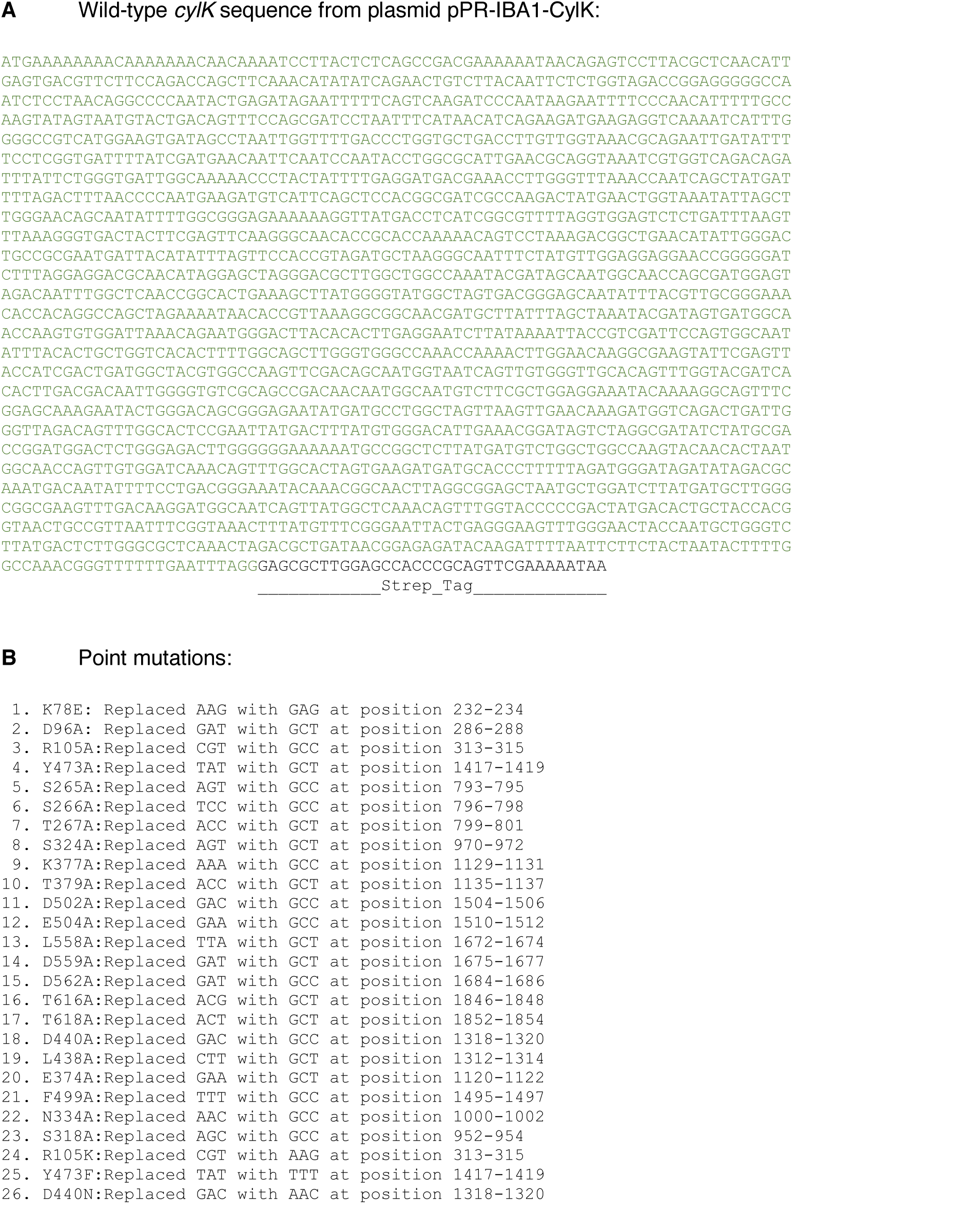
*(A)* Wild-type *cylK* DNA sequence used for soluble expression in activity assays. *(B)* Point mutations constructed and sequence verified by Genewiz (South Plainfield, NJ, USA).

**Figure S15.**
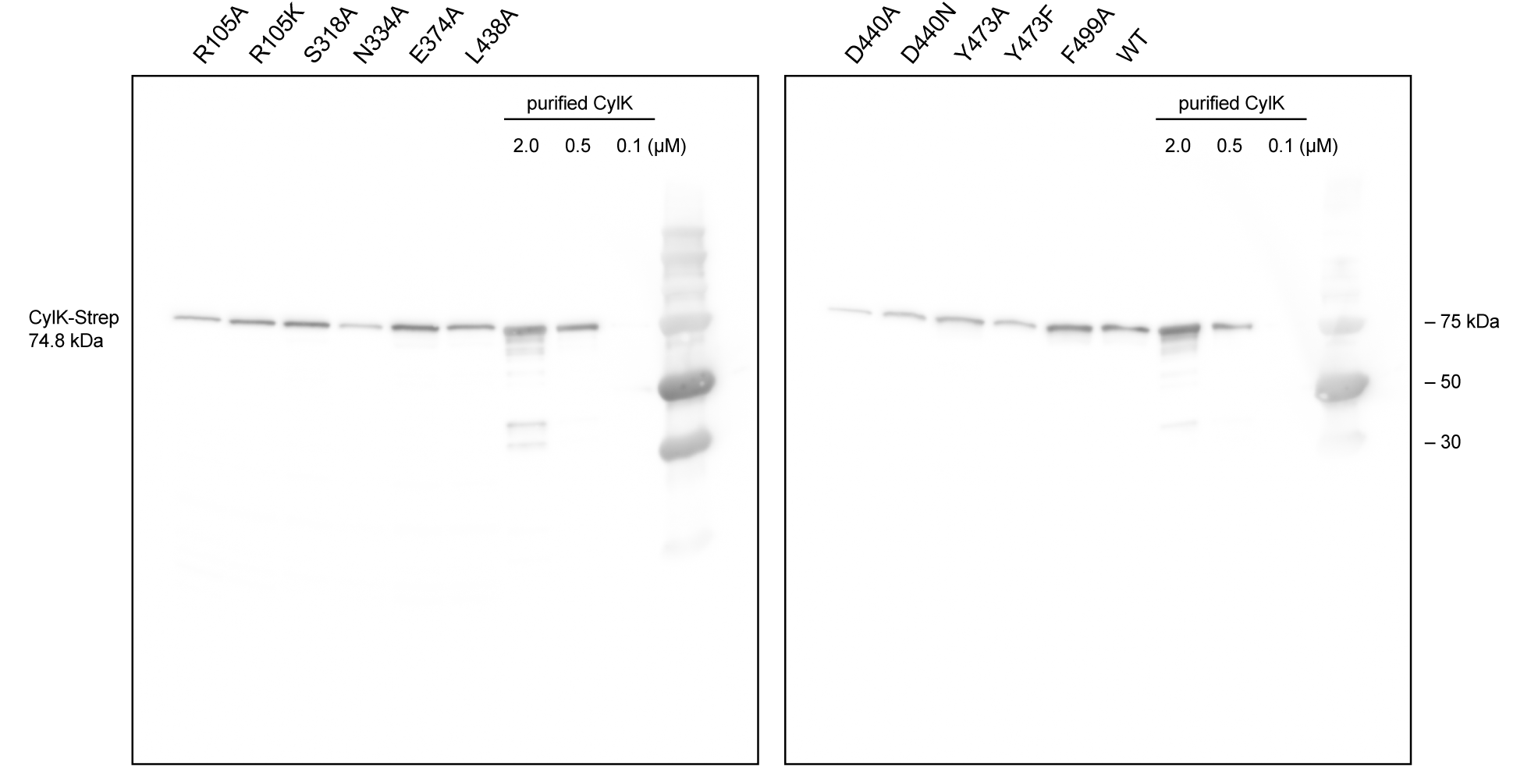
Western blotting of Strep-tagged CylK mutants from soluble lysate with anti-Strep-HRP (horseradish peroxidase, IBA) in order to ensure mutant enzyme expression and solubility. Serial dilution of purified CylK-Strep included to visualize protein concentration range.

**Table S1.** Data collection and refinement statistics for Tb-soaked *C. licheniforme* CylK structures.

|  | Tb anomalous | Native (final model) |
| --- | --- | --- |
| **Data collection** |  |  |
| Space group | *C* 2 2 2<sub>1</sub> | *C* 2 2 2<sub>1</sub> |
| Wavelength (Å) | 1.6314 | 0.97872 |
| Cell dimensions |  |  |
| *a*, *b*, *c* (Å) | 94.265, 139.286, 100.381 | 93.667, 138.613, 99.438 |
| $\alpha$, $\beta$, $\gamma$ (°) | 90.0, 90.0, 90.0 | 90.0, 90.0, 90.0 |
| Resolution (Å) | 50-2.38 (2.42-2.38) | 50.0-1.68 (1.71-1.68) |
| *R*<sub>merge</sub> | 0.218 (0.670) | 0.101 (1.326) |
| *R*<sub>pim</sub> | 0.041 (0.192) | 0.026 (0.346) |
| *I* / $\sigma I$ | 25.2 (3.8) | 29.3 (2.15) |
| CC<sub>1/2</sub> | 1.010 (0.906) | (0.780) |
| Completeness (%) | 99.5 (99.9) | 99.8 (99.3) |
| Redundancy | 29.5 (11.4) | 14.8 (14.3) |
| **Refinement** |  |  |
| Resolution (Å) |  | 1.68 |
| No. reflections |  | 68,476 |
| *R*<sub>work</sub> / *R*<sub>free</sub> |  | 0.1636/0.1980 |
| No. atoms |  | 5292 |
| Protein |  | 4903 |
| Ligand/ion |  | 50 |
| Water |  | 339 |
| *B*-factors |  |  |
| Protein |  | 18.04 |
| Ligand/ion |  | 27.97 |
| Water |  | 25.65 |
| R.m.s. deviations |  |  |
| Bond lengths (Å) |  | 0.012 |
| Bond angles (°) |  | 1.677 |
| Molprobability clashscore |  | 1.56 (99<sup>th</sup> percentile) |
| Rotamer outliers (%) |  | 1.36 |
| Ramachandran favored (%) |  | 94.77 |
| PDB accession code |  | 7RON |

\*Values in parentheses are for highest-resolution shell.

**Table S2.**
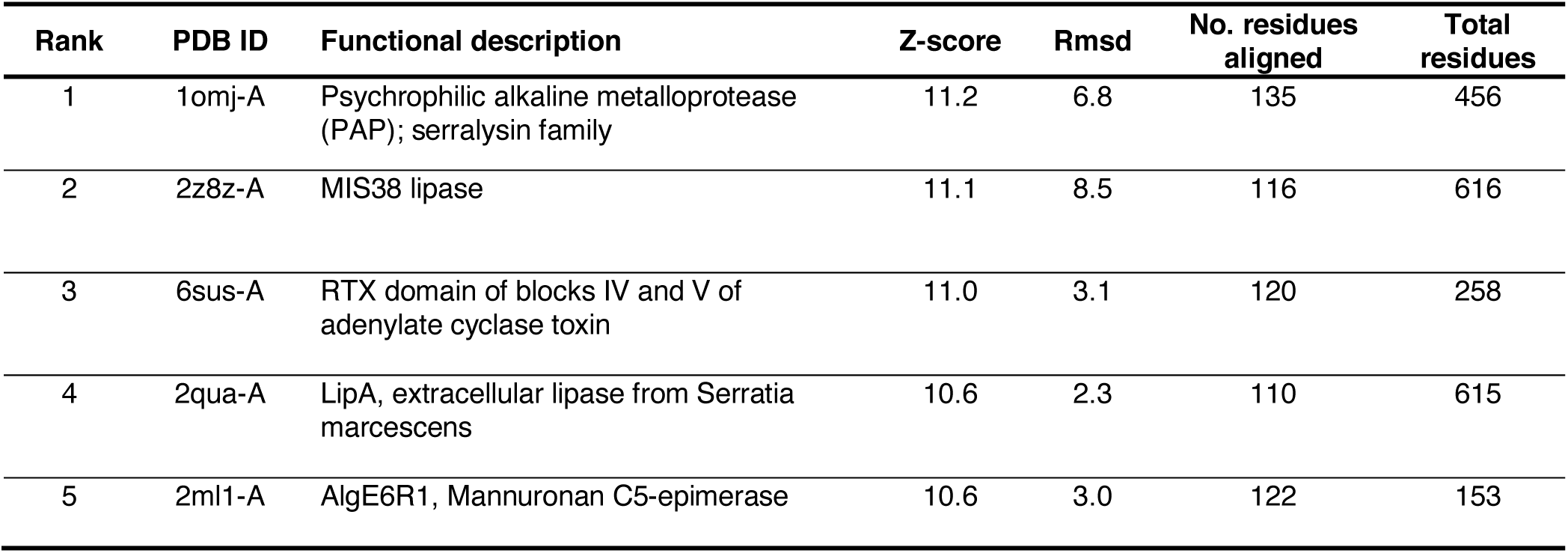
Selected structural homologs of the N-terminal domain of CylK obtained by a structural comparison to other proteins in the PDB (4).

| Rank | PDB ID | Functional description | Z-score | Rmsd | No. residues aligned | Total residues |
| --- | --- | --- | --- | --- | --- | --- |
| 1 | 1omj-A | Psychrophilic alkaline metalloprotease (PAP); serralyisin family | 11.2 | 6.8 | 135 | 456 |
| 2 | 2z8z-A | MIS38 lipase | 11.1 | 8.5 | 116 | 616 |
| 3 | 6sus-A | RTX domain of blocks IV and V of adenylate cyclase toxin | 11.0 | 3.1 | 120 | 258 |
| 4 | 2qua-A | LipA, extracellular lipase from *Serratia marcescens* | 10.6 | 2.3 | 110 | 615 |
| 5 | 2ml1-A | AlgE6R1, Mannuronan C5-epimerase | 10.6 | 3.0 | 122 | 153 |

**Table S3.** Selected structural homologs of the C-terminal domain of CylK obtained by a structural comparison to other proteins in the PDB (4).

| Rank | PDB ID | Functional description | Z-score | Rmsd | No. residues aligned | Total residues |
| --- | --- | --- | --- | --- | --- | --- |
| 1 | 1SQ9-A | Antiviral protein Ski8 | 31.2 | 2.4 | 286 | 378 |
| 2 | 4H5j-B | Guanine nucleotide-exchange factor Sec12 | 31.1 | 2.9 | 300 | 347 |
| 3 | 5TF2-A | Prolactin regulatory element-binding protein | 31.0 | 2.7 | 291 | 338 |
| 4 | 4J0X-B | Ribosomal RNA-processing protein 9 | 30.9 | 2.4 | 290 | 366 |
| 5 | 2Z2O-C | Virginiamycin B lyase | 30.5 | 2.4 | 286 | 299 |

**Table S4.** Data collection and refinement statistics for Br-soaked *C. licheniforme* CylK structures.

|  | Native | Br anomalous |
| --- | --- | --- |
| **Data collection** |  |  |
| Space group | *C* 2 2 2<sub>1</sub> | *C* 2 2 2<sub>1</sub> |
| Wavelength (Å) | 0.97872 | 0.9184 |
| Cell dimensions |  |  |
| *a*, *b*, *c* (Å) | 93.470, 138.656, 99.511 | 93.557, 138.726, 99.706 |
| $\alpha$, $\beta$, $\gamma$ (°) | 90.0, 90.0, 90.0 | 90.0, 90.0, 90.0 |
| Resolution (Å) | 50.0-1.52 (1.55-1.52) | 50-1.54 (1.57-1.54) |
| *R*<sub>merge</sub> | 0.076 (0.703) | 0.725 (4.755) |
| *R*<sub>pim</sub> | 0.029 (0.295) | 0.149 (1.140) |
| *I* / σ*I* | 23.3 (2.14) | 24.0 (2.06) |
| CC<sub>1/2</sub> | 0.990 (0.843) | 0.978 (0.124) |
| Completeness (%) | 99.7 (98.9) | 100.0 (100.0) |
| Redundancy | 7.0 (5.5) | 24.2 (16.9) |
| **Refinement** |  |  |
| Resolution (Å) | 1.52 |  |
| No. reflections | 89,122 |  |
| *R*<sub>work</sub> / *R*<sub>free</sub> | 0.1693/0.1978 |  |
| No. atoms | 5,437 |  |
| Protein | 4,989 |  |
| Ligand/ion | 45 |  |
| Water | 403 |  |
| *B*-factors |  |  |
| Protein | 16.68 |  |
| Ligand/ion | 23.03 |  |
| Water | 23.77 |  |
| R.m.s. deviations |  |  |
| Bond lengths (Å) | 0.013 |  |
| Bond angles (°) | 1.823 |  |
| Molprobability clashscore | 1.64 (99<sup>th</sup> percentile) |  |
| Rotamer outliers (%) | 0.76 |  |
| Ramachandran favored (%) | 93.90 |  |
| PDB accession code | 7ROO |  |

\*Values in parentheses are for highest-resolution shell.

